# PICH promotes SUMOylated TopoisomeraseIIα dissociation from mitotic centromeres for proper chromosome segregation

**DOI:** 10.1101/781401

**Authors:** Victoria Hassebroek, Hyewon Park, Nootan Pandey, Brooklyn T. Lerbakken, Vasilisa Aksenova, Alexei Arnaoutov, Mary Dasso, Yoshiaki Azuma

## Abstract

Polo-like kinase interacting checkpoint helicase (PICH) is a SNF2 family DNA translocase and is a Small Ubiquitin-like modifier (SUMO) binding protein. Despite that both translocase activity and SUMO-binding activity are required for proper chromosome segregation, how these two activities function to mediate chromosome segregation remains unknown. Here, we show that PICH specifically promotes dissociation of SUMOylated TopoisomeraseIIα (TopoIIα) from mitotic chromosomes. When TopoIIα is stalled by treatment of cells with a potent TopoII inhibitor, ICRF-193, TopoIIα becomes SUMOylated, and this promotes its interaction with PICH. Conditional depletion of PICH using the Auxin Inducible Degron (AID) system resulted in retention of SUMOylated TopoIIα on chromosomes, indicating that PICH removes stalled SUMOylated TopoIIα from chromosomes. *In vitro* assays showed that PICH specifically regulates SUMOylated TopoIIα activity using its SUMO-binding and translocase activities. Taken together, we propose a novel mechanism for how PICH acts on stalled SUMOylated TopoIIα for proper chromosome segregation.

**Summary Statement:** Polo-like kinase interacting checkpoint helicase (PICH) interacts with SUMOylated proteins to mediate proper chromosome segregation during mitosis. The results demonstrate that PICH promotes dissociation of SUMOylated TopoisomeraseIIα from chromosomes and that function leads to proper chromosome segregation.

## Introduction

Accurate chromosome segregation is a complex and highly regulated process during mitosis. Sister chromatid cohesion is necessary for proper chromosome alignment, and is mediated by both Cohesin and catenated DNA at centromeric regions (Bauer et al., 2012, Losada et al., 1998, Michaelis et al., 1997). Compared to the well-described regulation of Cohesin (Morales and Losada, 2018), the regulation of catenated DNA cleavage by DNA TopoisomeraseIIα (TopoIIα) is not fully understood despite its critical role in sister chromatid disjunction. ATP-dependent DNA decatenation by TopoIIα takes place during the metaphase-to-anaphase transition allowing for proper sister chromatid disjunction (Gomez et al., 2014, Shamu and Murray, 1992, Wang et al., 2010). Failure in resolution of catenanes by TopoIIα leads to the formation of chromosome bridges, and ultra-fine DNA bridges (UFBs) to which PICH localizes (Spence et al., 2007). PICH is a SNF2 family DNA translocase (Baumann et al., 2007, Biebricher et al., 2013), and its binding to UFBs recruits other proteins to UFBs (Chan et al., 2007, Hengeveld et al., 2015). In addition to the role in UFB binding during anaphase, PICH has been shown to play a key role in sister chromatid disjunction in the metaphase to anaphase transition (Baumann et al., 2007, Nielsen et al., 2015, Sridharan and Azuma, 2016). These studies suggest that PICH surveys for and resolves catenanes during prometaphase to metaphase, assuring the proper segregation of sister chromatids during anaphase.

Recently, we demonstrated that both DNA translocase activity and SUMO-binding activity of PICH are required for chromosome segregation (Sridharan and Azuma, 2016). PICH binds SUMOylated proteins using its three SUMO interacting motifs (SIMs) (Sridharan et al., 2015). PICH utilizes ATPase activity to translocate DNA similar to known nucleosome remodeling enzymes (Whitehouse et al., 2003), thus it is a putative remodeling enzyme for chromatin proteins. Intriguingly, the nucleosome remodeling activity of PICH was shown to be limited as compared to established nucleosome remodeling factors (Ke et al., 2011). Therefore, the target of PICH remodeling activity has not yet been determined. Importantly, both loss of function PICH mutants in either SUMO-binding activity or translocase activity showed chromosome bridge formation (Sridharan and Azuma, 2016), suggesting that both of these activities cooperate to accomplish proper chromosome segregation. Previous studies demonstrated that PICH-depleted cells have increased sensitivity to ICRF-193, a potent TopoII catalytic inhibitor, accompanied with increased incidence of chromosome bridges, binucleation, and micronuclei formation (Kurasawa and Yu-Lee, 2010, Nielsen et al., 2015, Wang et al., 2008). ICRF-193 stalls TopoIIα at the last step of the strand passage reaction (SPR) in which two DNA strands are trapped within TopoIIα without DNA strand breaks (Patel et al., 2000, Roca et al., 1994). Therefore, ICRF-193 treatment in mitotic cells produces unresolved catenanes bound by stalled TopoIIα. Recent studies indicate that PICH prevents this event by increasing TopoIIα activity (Nielsen et al., 2015). However, it is unknown how PICH resolves stalled TopoIIα in closed clamp conformation with ICRF-193 treatment. Importantly, it has been shown that ICRF-193 treatment increases SUMOylation of TopoIIα on mitotic chromosomes (Agostinho et al., 2008). This upregulation of TopoIIα SUMOylation was not observed after treatment with another potent TopoII inhibitor, Merbarone (Agostinho et al., 2008). Merbarone prevents TopoII activity at the initial stage of its SPR and in a different conformation than ICRF-193 (Fortune and Osheroff, 1998). Therefore, it does not result in stalled TopoII on DNA. This distinction between TopoII inhibitors suggests that SUMOylation of TopoIIα represents the stalled TopoIIα on DNA. These observations lead to the hypothesis that PICH interacts with SUMOylated TopoIIα and prevents the formation of chromosome bridges by resolving stalled TopoIIα-mediated catenanes.

Our results demonstrate that TopoIIα is SUMOylated upon stalling by ICRF-193 treatment, leading to recruitment of PICH to SUMOylated TopoIIα. Depletion of PICH retained more SUMOylated TopoIIα on the chromosomes in ICRF-193 treated cells. This suggests that PICH is required for removing stalled SUMOylated TopoIIα from chromosomes. *In vitro* assays suggest that PICH controls SUMOylated TopoIIα activity in both a translocase and SUMO-binding dependent manner. Together, we propose a novel mechanism for PICH in promoting proper chromosome segregation in mitosis by removing stalled SUMOylated TopoIIα from mitotic chromosomes.

## Results

### PICH, SUMO2/3, and TopoIIα colocalize on mitotic chromosomes upon TopoIIα inhibition by ICRF-193

Treatment with ICRF-193, a catalytic inhibitor of TopoII which blocks TopoII at the last stage of its SPR, after DNA decatenation but before DNA release, increases SUMO2/3 modification of TopoIIα on mitotic chromosomes. In contrast, treatment with another catalytic TopoII inhibitor, Merbarone, which blocks TopoII before the cleavage step of the SPR, does not affect the level of SUMO2/3 modification of TopoIIα (Agostinho et al., 2008). We utilized these two contrasting inhibitors to assess whether TopoIIα inhibition and/or SUMOylation changes PICH distribution on mitotic chromosomes. HCT116 cells were synchronized in prometaphase, and mitotic cells were isolated by shake off. To assess the effects of the TopoII inhibitors specifically during mitosis, the inhibitors were added to the cells after mitotic shake off. Consistent with previous reports (Agostinho et al., 2008), Western blotting analysis of isolated chromosomes showed that treatment with ICRF-193 increased the overall SUMO2/3 modification of chromosomal proteins. There were upshifted bands detected by anti-TopoIIα antibody, which indicate that the ICRF-193 treatment increased the level of SUMOylated TopoIIα on chromosomes (marked by red asterisks in Figure 1A). In contrast, Merbarone did not induce apparent differences in either the overall SUMO2/3 modification of chromosomal proteins or the TopoIIα SUMOylation, suggesting the specificity of ICRF-193 on the SUMOylation of TopoIIα (Figure 1A).

**Figure 1.**
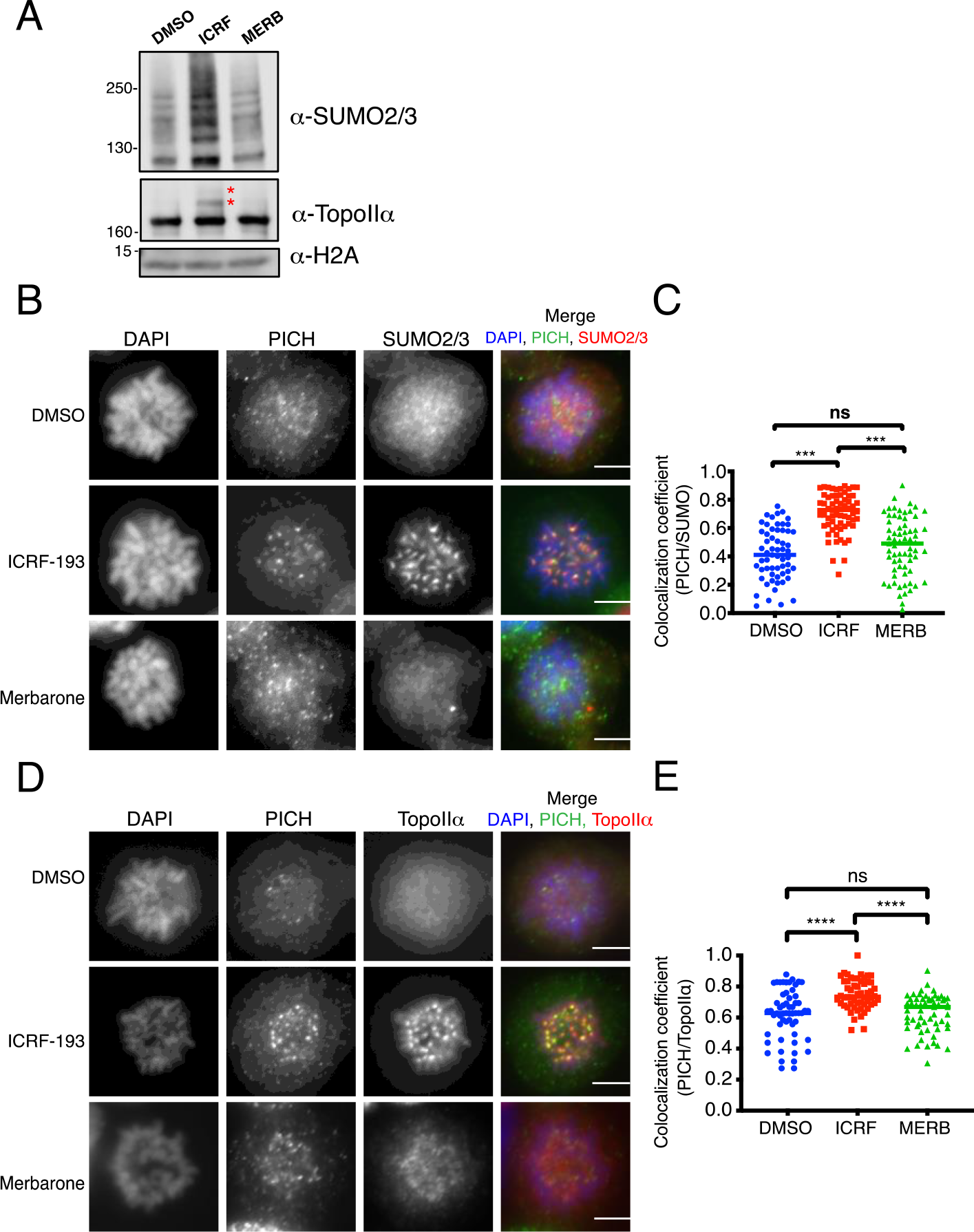
TopoIIα inhibition by ICRF-193 leads to PICH/SUMO2/3 and PICH/TopoIIα colocalization. **(A)** HCT116 cells were synchronized and treated with indicated inhibitors (ICRF-193: ICRF, and Merbarone: Merb). Mitotic chromosomes were isolated and subjected to Western blotting with indicated antibodies. * indicates SUMOylated TopoIIα. **(B)** Mitotic cells treated with DMSO, ICRF-193, and Merbarone were stained with antibodies against: PICH (green) and SUMO2/3 (red). DAPI shows DNA (blue). Scale bar = 10μm. **(C)** A minimum of twenty prometaphase cell images were obtained in each group from three independent experiments, and the colocalization coefficients between PICH/SUMO2/3 was calculated and plotted. **(D)** Mitotic cells were treated as in **B** and stained with antibodies against: PICH (green), TopoIIα (red). DNA was stained with DAPI (blue). Scale bar = 10μm. **(E)** Colocalization coefficients were analyzed as described in **C**. p values for comparison among three experiments were calculated using a one-way ANOVA analysis of variance with Tukey multi-comparison correction. ns: not significant; *: p ≤ 0.05; ***: p < 0.001; ****: p < 0.0001

Immunofluorescent analysis of cells treated with ICRF-193 or Merbarone showed SUMO 2/3 foci at the centromere consistent with previous studies (Azuma et al., 2005; Zhang et al., 2008). However, in ICRF-193 treated cells there was an increase in SUMO2/3 foci intensity as compared to cells treated with DMSO or Merbarone (Figure 1B). Similar to previous reports, PICH foci were observed at the centromere (Baumann et al., 2007, Sridharan and Azuma, 2016). Again, PICH foci intensity was increased in cells treated with ICRF-193. Then, the colocalization of PICH and SUMO2/3 was measured in the entire cell. The incidence of colocalization between PICH and SUMO2/3 signals in the ICRF-193 treated cells was significantly higher (72% on average) than DMSO treated cells (42% on average) (Figure 1B middle row, C). In contrast, the incidence of colocalization between SUMO2/3 and PICH in the Merbarone treated cells did not show any significant changes compared to DMSO treated cells (Figure 1B bottom row, C).

Because our previous study showed that PICH binds to SUMOylated TopoIIα C-terminal domain (Sridharan et al., 2015), we anticipated that the increased incidence of colocalization between SUMO2/3 and PICH is due to the interaction between these two molecules. When cells were treated with ICRF-193, the incidence of colocalization between PICH and TopoIIα at the centromere was significantly higher than the DMSO treated cells (Figure 1D, E). In contrast, Merbarone treatment did not show any significant differences as compared to DMSO treated cells (Figure 1D, E). This suggests that ICRF-193 mediated SUMOylation promotes a redistribution of PICH resulting in both PICH/TopoIIα and PICH/SUMO2/3 colocalization at centromeric regions. Because the colocalization between PICH/SUMO2/3 and PICH/TopoIIα is induced by ICRF-193 but not by Merbarone, the colocalization of these proteins is triggered by the hyper-accumulation of SUMOylated proteins at the centromere but not by inhibition of TopoIIα activity.

### SUMOylated TopoIIα is a critical binding target of PICH in ICRF-193 treated cells

To determine if SUMOylated TopoIIα is the target of PICH in ICRF-193 treated cells, we examined whether TopoIIα depletion (ΔTopoIIα) affects PICH/SUMO2/3 colocalization. To deplete TopoIIα from cells, we created a conditional TopoIIα-knockdown cell line which utilizes the Auxin-Inducible Degron (AID) system (Natsume et al., 2016, Nishimura et al., 2009) (Supplemental Figure S1 and S2). We inserted DNA encoding an AID-Flag tag into the TopoIIα locus (Supplemental Figure S2A, B, and C) to a cell line that has verified integration of the *OsTIR1* gene, an auxin-dependent Ubiquitin E3 ligase, in the genome (Supplemental Figure S1). After auxin addition AID-tagged TopoIIα was degraded to undetectable levels within 6 hours. (Supplemental Figure S2D). This rapid elimination allows us to examine the effect of TopoIIα depletion in a single cell cycle. In ΔTopoIIα ICRF-193 treated cells, there was an extreme reduction of SUMO2/3 and PICH signals at centromeres (marked by CENP-C) (Figure 2A). In addition, the incidence of colocalization between PICH/SUMO2/3 at the centromere was also significantly reduced in ΔTopoIIα ICRF-193-treated cells (Figure 2B comparing red and purple characters). This suggests that PICH is recruited to SUMOylated TopoIIα in the presence of ICRF-193. Consistently, the Western blotting analysis using isolated mitotic chromosomes obtained from ΔTopoIIα ICRF-193-treated cells showed slightly decreased SUMO2/3 modification and reduced level of PICH compared to control cells treated with ICRF-193 (Figure 2C +Auxin/ICRF lane). Although, because the levels of SUMO2/3 modification were still increased as compared to DMSO treated control cells, that may represent, in part, SUMOylation of TopoIIβ TopoIIβ has been shown to have increased SUMOylation under ICRF-193 treatment (Isik et al., 2003, Mao et al., 2000). Because PICH/SUMO2/3 colocalization is decreased to control levels when TopoIIα is depleted, TopoIIβ is clearly not the target to which PICH interacts. Together, the results show PICH targets SUMOylated TopoIIα at centromeres under ICRF-193 treatment.

**Figure 2.**
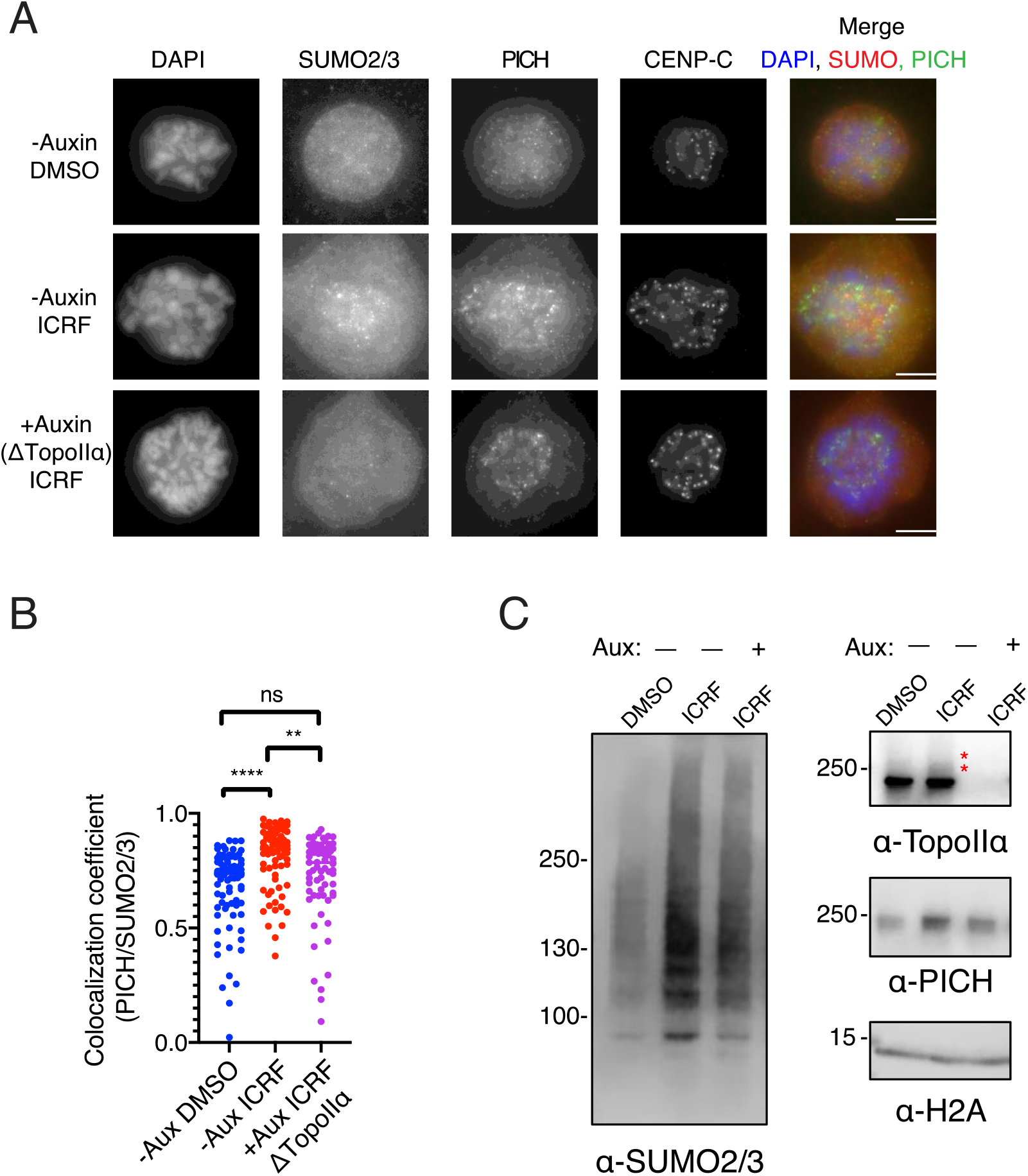
Depletion of TopoIIα attenuates SUMO2/3 modification and decreases PICH/SUMO2/3 colocalization at the centromere in ICRF-193 treated cells. **(A)** DLD-1 cells with endogenous TopoIIα tagged with an AID were synchronized in mitosis and treated with DMSO and ICRF-193. Auxin was added to the cells for 6 hours after Thymidine release. Mitotic cells were fixed and stained with antibodies against: SUMO2/3 (red), PICH (green), CENP-C (not merged). DNA was stained with DAPI (blue). Scale bar = 10μm. **(B)** The colocalization coefficients between PICH and SUMO2/3 of cells with indicated treatments (−/+Auxin and DMSO or ICRF) were measured. p values for comparison among four experiments were calculated using a one-way ANOVA analysis of variance with Tukey multi-comparison correction; ns: not significant; **: p < 0.01; ****: p < 0.0001. **(C)** Mitotic chromosomes were isolated with (+Aux) or without (-Aux) auxin treatment and DMSO or ICRF-193 and subjected to Western blotting with indicated antibodies. * indicates SUMOylated TopoIIα.

### SUMOylation is required for PICH/TopoIIα colocalization at the centromere

The increased PICH localization to centromeres in ICRF-193 treated cells is likely due to TopoIIα SUMOylation. To determine whether SUMOylation is required for PICH localization to mitotic centromeres, we established cell lines to attenuate the level of SUMOylation at the centromere. To accomplish this, we generated a fusion protein, called Py-S2, which consists of the SENP2-catalytic domain (required for deSUMOylation) (Reverter and Lima, 2004, Ryu et al., 2015, Sridharan et al., 2015), and of the N-terminal region of human PIASy (localizes to mitotic centromeres through its specific binding with the RZZ complex at the kinetochore) (Ryu and Azuma, 2010)). As a negative control, we substituted a cysteine at the position 548 of SENP2 to an alanine (called Py-S2 Mut) to create a loss of function mutant of the SENP2 deSUMOylation activity (Reverter and Lima, 2004, Reverter and Lima, 2006) (Figure 3A). The activity of the recombinant fusion proteins was verified by Xenopus egg extract (XEE) assay (Supplemental Figure S3). As we predicted, addition of the Py-S2 protein to XEE completely eliminates mitotic chromosomal SUMOylation. To our surprise, Py-S2 Mut protein stabilized the SUMOylation of chromosomal proteins, thus acted as a dominant negative mutant. To express the fusion proteins in cells, we created inducible expression cell lines using the Tetracycline inducible system (Supplemental Figure S4). We integrated each of the fusion genes into the AAVS1 safe harbor locus of HCT116 cells using integration plasmids (Natsume et al., 2016).

**Figure 3.**
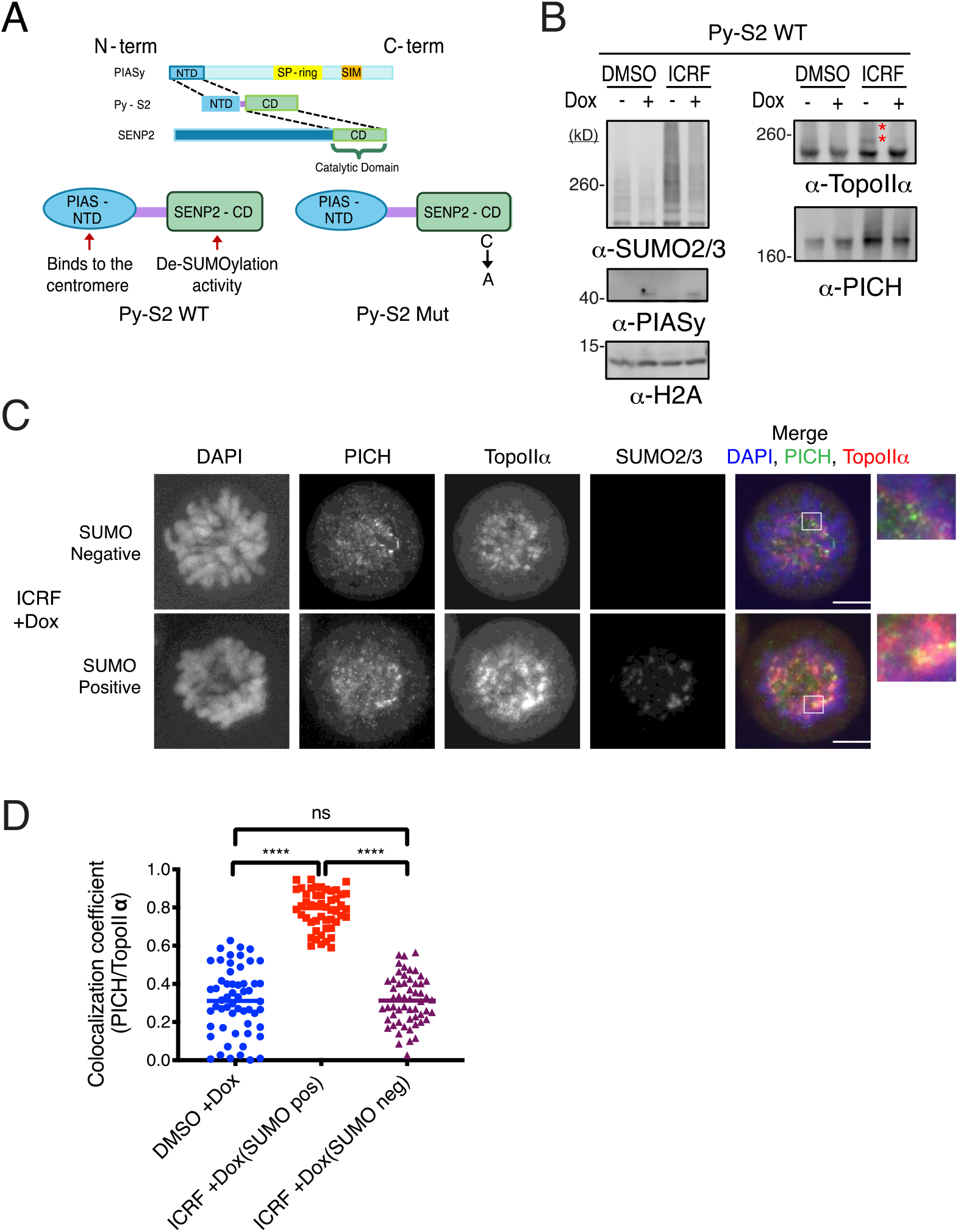
DeSUMOylation enzyme inhibits PICH/TopoIIα colocalization on chromosomes. **(A)** Schematic of fusion proteins generated for modulating SUMOylation on mitotic chromosomes. **(B)** Mitotic chromosomes were subjected to Western blotting with indicated antibodies. *indicates SUMOylated TopoIIα. **(C)** Mitotic cells were fixed and stained with antibodies against: PICH (green), TopoIIα (red), and SUMO2/3 (far red). DNA was stained by DAPI (blue). Scale bar = 10μm. **(D)** The colocalization coefficients between PICH and TopoIIα were measured in the cells with indicated treatments (+Doxycycline, +ICRF-193 or +DMSO) and categories (SUMO pos or SUMO neg). p values for comparison among three experiments were calculated using a one-way ANOVA analysis of variance with Tukey multi-comparison correction. ns: not significant; ****: p < 0.0001

To test the effect of attenuated SUMOylation during mitosis, cells were synchronized with or without doxycycline treatment, and chromosomes were isolated. Western blotting of mitotic chromosomal fractions showed that expression of Py-S2 attenuated the SUMO2/3 modification on mitotic chromosomes. The attenuation of mitotic SUMO2/3 modification by Py-S2 became apparent in the ICRF-193 treated samples (Figure 3B comparing −/+Dox with ICRF-193). Consistent with the SUMO2/3 modification profile, Py-S2 expression substantially attenuated SUMOylated TopoIIα in ICRF-193 treated cells (Figure 3B comparing −/+Dox samples with ICRF-193). Notably, the amount of PICH on chromosomes in the ICRF-193 treated sample was reduced when SUMOylation was attenuated by Py-S2 (Figure 3B comparing −/+Dox samples with ICRF-193). Distinct from the result that displayed complete elimination of SUMO2/3 modification in XEE assays (Supplemental Figure S3A), the SUMO2/3 and SUMOylated TopoIIα signals were still present in the Py-S2 expressing cells. This is likely due to the mosaic expression of Py-S2 represented by SUMO2/3 signals (Figure 3C). The SUMO2/3 signals in Py-S2 expressing cells displayed either retention of SUMO2/3 signals on chromosomes suggesting no transgene expression (hereafter referred to as SUMO positive cells), or no SUMO2/3 signal on chromosomes suggesting transgene expression (hereafter referred to as SUMO negative cells) (Figure 3C). PICH signals in the SUMO positive cells displayed strong centromeric foci, resembling that of ICRF-193-treated parental HCT116 cells (Figure 1B). In SUMO negative cells, the intensity of PICH foci was lessened and a more diffuse non-centromeric signal was observed. Consistent with our observations in XEE assays, TopoIIα signal did not show apparent differences between SUMO positive cells and negative cells, suggesting that inhibition of mitotic SUMOylation does not affect TopoIIα association with chromosomes (Azuma et al., 2005, Azuma et al., 2003). Colocalization between PICH and TopoIIα foci was reduced in SUMO negative cells (indicated by the magnified image in the merged panel in Figure 3C). The incidence of PICH/TopoIIα colocalization in the SUMO negative/ICRF-193 treated cells was comparable to DMSO treated cells (Figure 3D comparing blue and purple characters), indicating that mitotic SUMOylation is indispensable for the ICRF-193 effect on PICH/TopoIIα colocalization. In SUMO positive cells, ICRF-193 treatment showed similar PICH/TopoIIα colocalization to parental HCT116 cells (comparing Figure 3D blue and red characters to those in Figure 1E). This strongly suggests that SUMOylation of TopoIIα is required for ICRF-193 induced PICH/TopoIIα colocalization.

Consistent with the result obtained from XEE assay (Supplemental Figure S3), Western blotting of the mitotic chromosome fraction showed that Py-S2 Mut expression in the HCT116 increased both overall SUMO2/3 modification and TopoIIα SUMOylation (Figure 4A +Dox samples). In addition, the amount of PICH on chromosomes was also increased in the Py-S2 Mut expressing cells (Figure 4A comparing −/+Dox samples). The immunostaining of Py-S2 Mut expressing cells revealed increased intensity of PICH foci as compared to uninduced cells treated with DMSO (Figure 4B DMSO −/+Dox). ICRF-193 treated Py-S2 Mut expressing cells showed a large amplification of PICH signal (Figure 4B ICRF −/+Dox). Importantly, the cells expressing the Py-S2 Mut showed a significant increase in PICH/TopoIIα colocalization independent of ICRF-193 treatment (Figure 4C comparing +Dox with DMSO or ICRF-193, blue and red boxes). Indicating that ICRF-193 treatment with Py-S2 Mut expression did not show a synergistic effect on PICH/TopoIIα colocalization (Figure 4C −/+ Dox with ICRF-193, red boxes). This suggests that increased chromosomal SUMOylation, presumably TopoIIα SUMOylation, is the major cause of ICRF-193 mediated PICH/TopoIIα colocalization on mitotic chromosomes. Together, these results further strengthen the concept that PICH specifically targets SUMOylated TopoIIα under ICRF-193 treatment.

**Figure 4.**
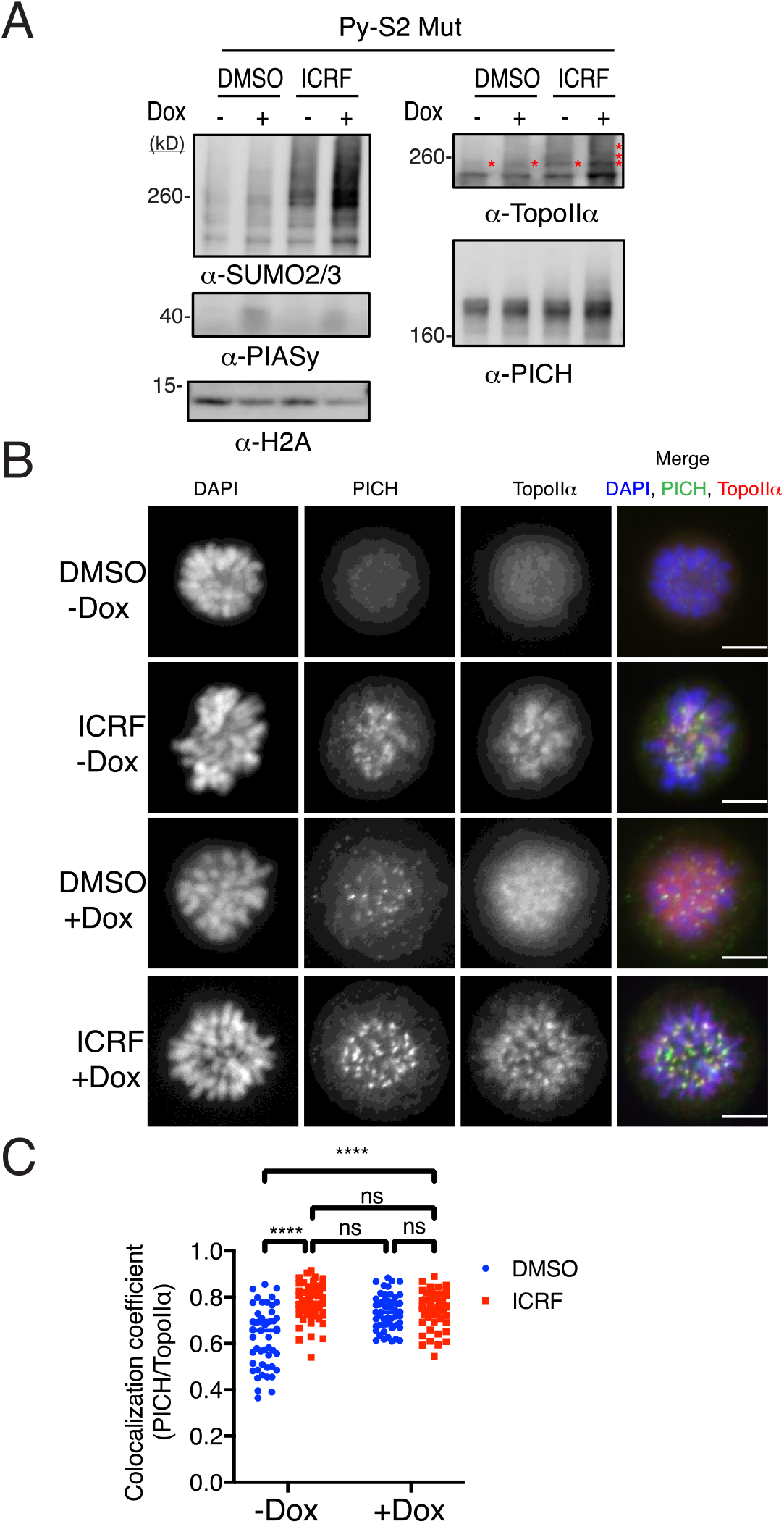
Dominant mutant of deSUMOylation enzyme promotes PICH/TopoIIα colocalization on chromosomes. **(A)** Mitotic chromosomes were isolated and subjected to Western blotting with indicated antibodies. *indicates SUMOylated TopoIIα. **(B)** Mitotic cells were treated as indicated and fixed then stained with antibodies against: PICH (green) and TopoIIα (red). DNA was stained with DAPI (blue). Scale bar = 10μm. **(C)** The colocalization coefficients between PICH and TopoIIα of cells with indicated treatments (+/− Doxycycline, +ICRF-193 or +DMSO) were measured. p values for comparison among three experiments were calculated using a two-way ANOVA analysis of variance with Tukey multi-comparison correction. ns: not significant; ****: p < 0.0001

### PICH controls the association of SUMOylated TopoIIα with chromosomes at the centromere

Our results suggest that PICH interacts with SUMOylated TopoIIα on mitotic chromosomes in ICRF-193 treated cells. To examine the biological function of PICH targeting SUMOylated TopoIIα, we generated a conditional PICH-knockdown cell using the AID system. DNA encoding an AID-Flag tag was introduced at the PICH locus (Supplemental Figure S5A, B, and C) into the OsTIR1-expressing stable cell line. Western blotting analysis confirmed that after auxin treatment AID-tagged PICH can be degraded to undetectable levels within 4 hours (Supplemental Figure S5D). To examine the effect of PICH depletion (ΔPICH) on SUMOylated TopoIIα, cells were synchronized in mitosis and chromosomes were isolated. Western blotting analysis of mitotic chromosomes showed that treatment with ICRF-193 increases the amount of PICH (Figure 5A -Auxin DMSO/ICRF lanes). This is consistent with results from native PICH expressing cells (Figure 2C, 3B, and 4B), suggesting that tagging PICH with the AID did not alter its response to ICRF-193 treatment. With the addition of auxin, there was no detectable PICH remaining on mitotic chromosomes (Figure 5A +Auxin DMSO/ICRF lanes). Importantly, PICH-depletion did not affect the amount of non-SUMOylated TopoIIα (Figure 5B +Auxin ΔPICH) but retained significantly higher levels of SUMOylated TopoIIα on mitotic chromosomes when the cells were treated with ICRF-193 (Figure 5C +Auxin ΔPICH). This suggests that PICH attenuates the interaction of SUMOylated TopoIIα with chromosomes. The function of PICH is amplified by the treatment of cells with ICRF-193, presumably due to increased levels of SUMOylated TopoIIα.

**Figure 5.**
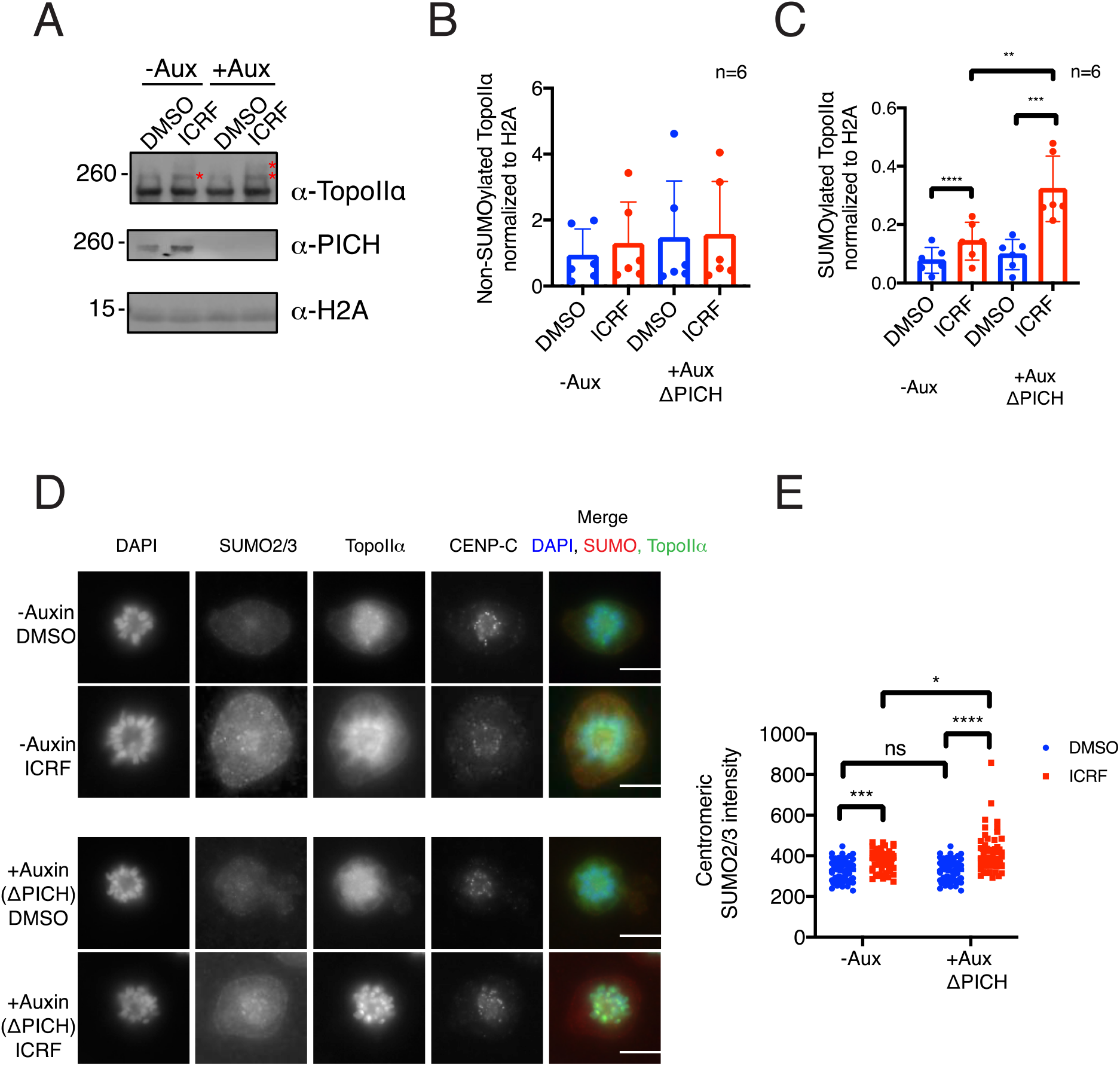
Chromosomes of PICH depleted cell show increased level of SUMOylated TopoIIα. **(A)** DLD-1 cells with endogenous PICH tagged with an AID were synchronized in mitosis and treated with DMSO and ICRF-193. Auxin was added to the cells for 6 hours after Thymidine release. Mitotic chromosomes were isolated and subjected to Western blotting with indicated antibodies. *indicates SUMOylated TopoIIα. **(B)** The intensity of non-SUMOylated TopoIIα signals normalized to H2A loading control. **(C)** The intensity of SUMOylated TopoIIα signals normalized to H2A loading control. **(D)** Mitotic cells were fixed and stained with antibodies against: SUMO2/3 (red), TopoIIα (green), CENP-C (far red). DNA was stained with DAPI (blue). Scale bar = 10μm. **(E)** The intensity of centromeric SUMO2/3 from four independent experiments was measured using FIJI software. Statistical analysis of **B** and **C** and **E** were performed by using a one-way ANOVA analysis of variance with Tukey multi-comparison correction; p values for comparison among four conditions were calculated. ns: not significant; **: p < 0.01; ***: p < 0.001; ****: p < 0.0001.

Consistent with Western blotting analysis, immunostaining of auxin treated cells showed no PICH signal on mitotic chromosomes in both DMSO and ICRF-193 treated cells (Supplemental Figure 6A +Auxin ΔPICH). The elimination of PICH foci on mitotic chromosomes was observed in all of the analyzed cells. There was no difference in TopoIIα localization in DMSO-treated cells with or without PICH, both showing diffuse signals on chromosomes with enrichment at the centromeres. This suggests that ΔPICH does not affect global TopoIIα localization in this analysis (Figure 5D comparing top row and third row). With ICRF-193 treatment, both PICH-nondepleted and ΔPICH cells showed further enrichment of TopoIIα foci at the centromere (Figure 5D comparing second row and bottom row). But there was less diffuse TopoIIα signal in ΔPICH cells treated with ICRF-193. The intensities of SUMO2/3 foci at centromeres were clearly increased in ΔPICH cells treated with ICRF-193 (Figure 5D bottom row). The quantification of centromeric SUMO2/3 foci indicated a statistically significant increase in the ΔPICH treated with ICRF-193 (Figure 5E). Because centromeric SUMO2/3 signal in ICRF-193 treated ΔTopoIIα cells was diminished (Figure 2A) and ΔPICH increased retention of SUMOylated TopoIIα on chromosomes (Figure 5C), the increased centromeric SUMO2/3 foci in ΔPICH ICRF-193 treated cells likely represent SUMOylated TopoIIα molecules. Together, these data indicate that PICH functions to remove stalled SUMOylated TopoIIα from mitotic centromeres in ICRF-193 treated cells.

**Figure 6.**
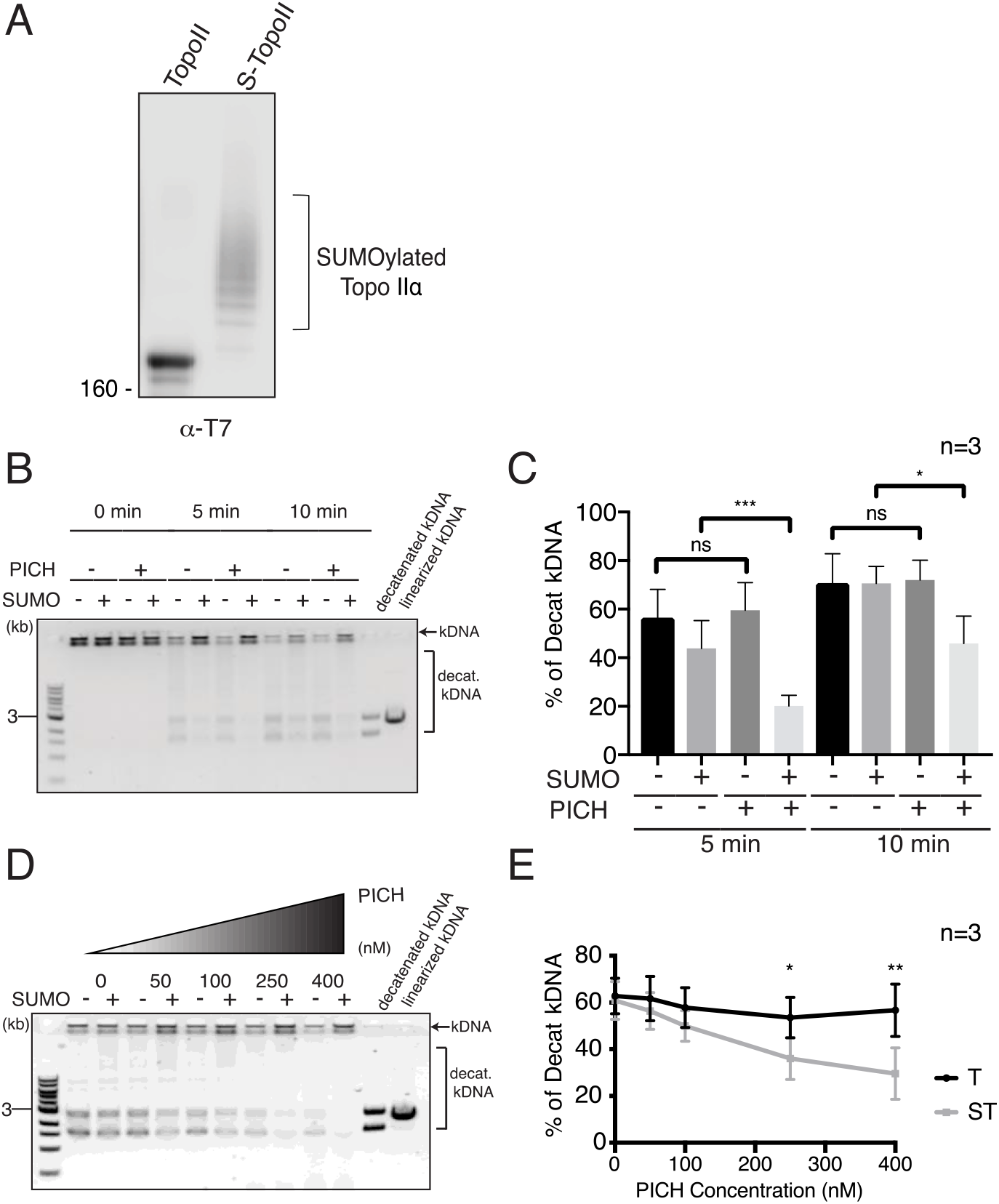
PICH inhibits SUMOylated TopoIIα decatenation activity. **(A)** Recombinant T7 tagged TopoIIα proteins were SUMOylated *in vitro*. Samples were subjected to Western blotting using anti-T7 tag antibody. **(B)** Decatenation of catenated kDNA in the indicated conditions (+/− PICH with non-SUMOylated TopoIIα (indicated as **—** SUMO) or SUMOylated TopoIIα (indicated as +SUMO)) were analyzed by DNA gel electrophoresis. Catenated kDNA is indicated by an arrow. Brackets indicate the decatenated kDNA species. **(C)** The decatenation activity of reactions in B was calculated as a percentage of decatenated kDNA. **(D)** Decatenation of catenated kDNA in SUMOylated and non-SUMOylated TopoIIα were analyzed by DNA gel electrophoresis with increasing concentrations of PICH. Catenated kDNA is indicated by an arrow. Brackets indicate decatenated kDNA species. **(E)** The decatenation activity of SUMOylated (ST) and non-SUMOylated TopoIIα (T) in D was calculated as a percentage of decatenated kDNA. Statistical analysis of **C** and **E** were performed by using a two-way ANOVA analysis of variance with Tukey multi-comparison correction; p values for comparison among six conditions were calculated. ns: not significant; *: p ≤ 0.05; **: p < 0.01; ***: p < 0.001

### Regulation of SUMOylated TopoIIα activity is dependent on both PICH ATPase activity and SIMs *in vitro*

Results from PICH-depleted cells suggest that PICH removes stalled SUMOylated TopoIIα induced by ICRF-193 from chromosomes. This activity may utilize both translocase activity of PICH and SUMO-binding activity to promote dissociation of SUMOylated TopoIIα from chromosomes. To examine if PICH SUMO-binding activity and translocase activity are important in controlling SUMOylated TopoIIα binding to DNA, we performed an *in vitro* DNA decatenation assay comparing non-SUMOylated and SUMOylated TopoIIα (Figure 6A) in the presence of PICH. Using the same conditions established in our previous study, recombinant *Xenopus laevis* TopoIIα was SUMOylated *in vitro*, then its DNA decatenation activity was analyzed by using catenated kDNA as the substrate (Ryu et al., 2010b). The decatenation activity was measured by calculating the percentage of decatenated kDNA separated by gel electrophoresis. On average, 70% of kDNA is decatenated at the ten-minute time-point when non-SUMOylated TopoIIα is present in the reaction (Figure 6B PICH **—** and SUMO **—** lanes). As we have previously shown, the decatenation activity of SUMOylated TopoIIα was reduced compared to non-SUMOylated TopoIIα. Importantly, when we added PICH to each of the reaction at concentrations equimolar to TopoIIα, the decatenation activity of SUMOylated TopoIIα was further attenuated (Figure 6B, C). The reduction of decatenation activity of SUMOylated TopoIIα was statistically significant at both the five minute and ten-minute time-points (Figure 6C light grey bars). Consistent with recent reports, the addition of PICH slightly increased decatenation activity of non-SUMOylated TopoIIα (Nielsen et al., 2015). A dose-dependent effect of PICH on SUMOylated TopoIIα decatenation activity was observed but that was not the case for non-SUMOylated TopoIIα. The concentration of TopoIIα in the reaction was 200nM, and PICH significantly reduced decatenation activity of SUMOylated TopoIIα ranging between 250nM up to 400nM (Figure 6D, E). Only SUMOylated TopoIIα was inhibited by PICH dose-dependently which is distinct from the PICH/non-SUMOylated TopoIIα interaction.

Because the translocase activity of PICH removes proteins from DNA, PICH inhibits decatenation activity of SUMOylated TopoIIα by removing SUMOylated TopoIIα from kDNA. To gain insight into that potential mechanism, we utilized a PICH mutant that has defects in either the SUMO-binding activity (PICH-d3SIM) or in translocase activity (PICH-K128A) (Figure 7A) (Sridharan et al., 2016). If PICH/SUMO interaction is critical for inhibiting decatenation activity of SUMOylated TopoIIα, the PICH-d3SIM mutant would lose its inhibitory function. In addition, we also expect that the PICH translocase activity deficient (PICH-K128A) mutant loses the ability to inhibit the decatenation activity of SUMOylated TopoIIα, because the mutant could not remove SUMOylated TopoIIα from kDNA. Supporting our hypothesis, PICH-d3SIM inhibited SUMOylated TopoIIα decatenation activity substantially less than wild-type (Figure 7B, C), suggesting that the direct SUMO/SIM interactions between PICH and SUMOylated TopoIIα plays a key role in this inhibition. Intriguingly, the translocase deficient PICH mutant further suppressed the SUMOylated TopoIIα decatenation activity (Figure 7B, D). This suggests that the translocase deficient mutant forms a stable complex with SUMOylated TopoIIα and the catenated kDNA because the translocase mutant retains its DNA binding ability (Kaulich et al., 2012, Nielsen et al., 2015, Sridharan and Azuma, 2016). Notably, neither mutant showed any significant effect on non-SUMOylated TopoIIα (Figure 7B) similar to wild-type PICH. This suggests that PICH binding to DNA does not inhibit the decatenation activity of TopoIIα, but rather it forms a complex with SUMOylated TopoIIα and prevents its decatenation activity. Taken together, our results suggest that PICH recognizes the SUMO moieties on TopoIIα through its SIMs and removes SUMOylated TopoIIα from DNAs using its translocase activity.

**Figure 7.**
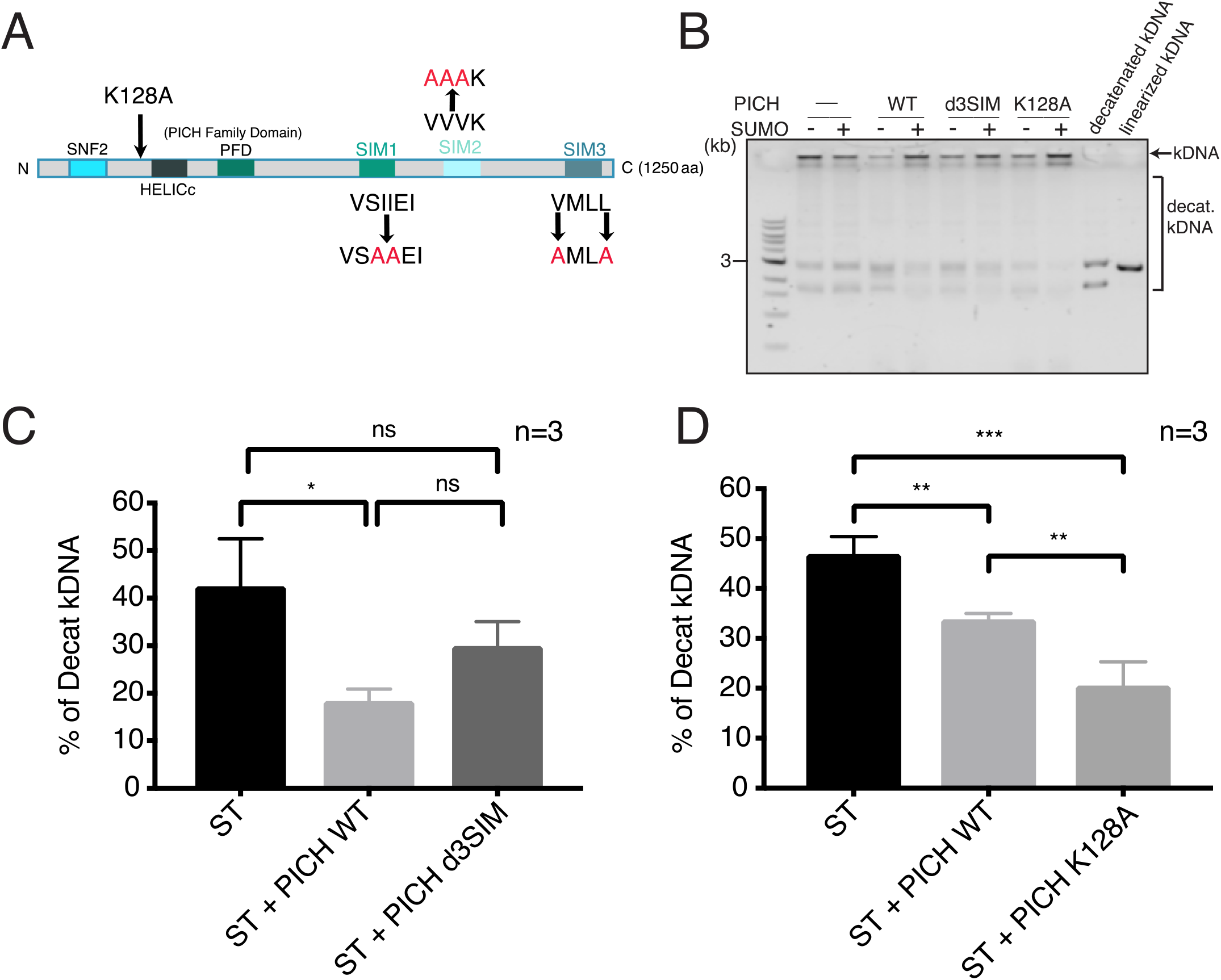
Both SUMO-binding activity and translocase activity of PICH involved in regulation of SUMOylated TopoIIα decatenation activity. **(A)** Schematic of PICH protein with known functional motifs. The introduced mutations in SIMs and in the ATPase domain (K128A) are indicated. **(B)** Representative gel showing non-SUMOylated (-SUMO) and SUMOylated TopoIIα (+SUMO) activity with PICH WT, a non-SUMO-binding mutant (d3SIM), and a translocase deficient mutant (K128A) or no PICH protein (-PICH). Catenated kDNA is indicated with an arrow. Brackets indicate decatenated kDNA species. **(C)** Decatenation activity of SUMOylated TopoIIα (ST) with indicated PICH (ST: no PICH, ST + PICH WT: PICH wild-type, ST + PICH d3SIM: PICH-d3SIM mutant). **(D)** Decatenation activity of SUMOylated TopoIIα (ST) with indicated PICH (ST: no PICH, ST + PICH WT: PICH wild-type, ST + PICH K128A: PICH-K128A mutant). Statistical analysis of **B** and **C** were performed by using a one-way ANOVA analysis of variance with Tukey multi-comparison correction; p values for comparison among six conditions were calculated. ns: not significant; *: p ≤ 0.05; **: p < 0.01; ***: p < 0.001

In conclusion, our results show that PICH targets SUMOylated TopoIIα to attenuate its interaction with chromosomes. When SUMOylation of TopoIIα is enhanced by its inhibitor, ICRF-193, the activity of PICH to remove SUMOylated TopoIIα from DNA becomes more prominent. Because ICRF-193 promotes trapped TopoIIα on DNA at the last stage of its SPR in a closed clamp conformation, we propose a model showing how PICH resolves detangled but trapped DNA that are bound within SUMOylated TopoIIα in both unperturbed and ICRF-193 affected mitosis (Figure 8).

**Figure 8.**
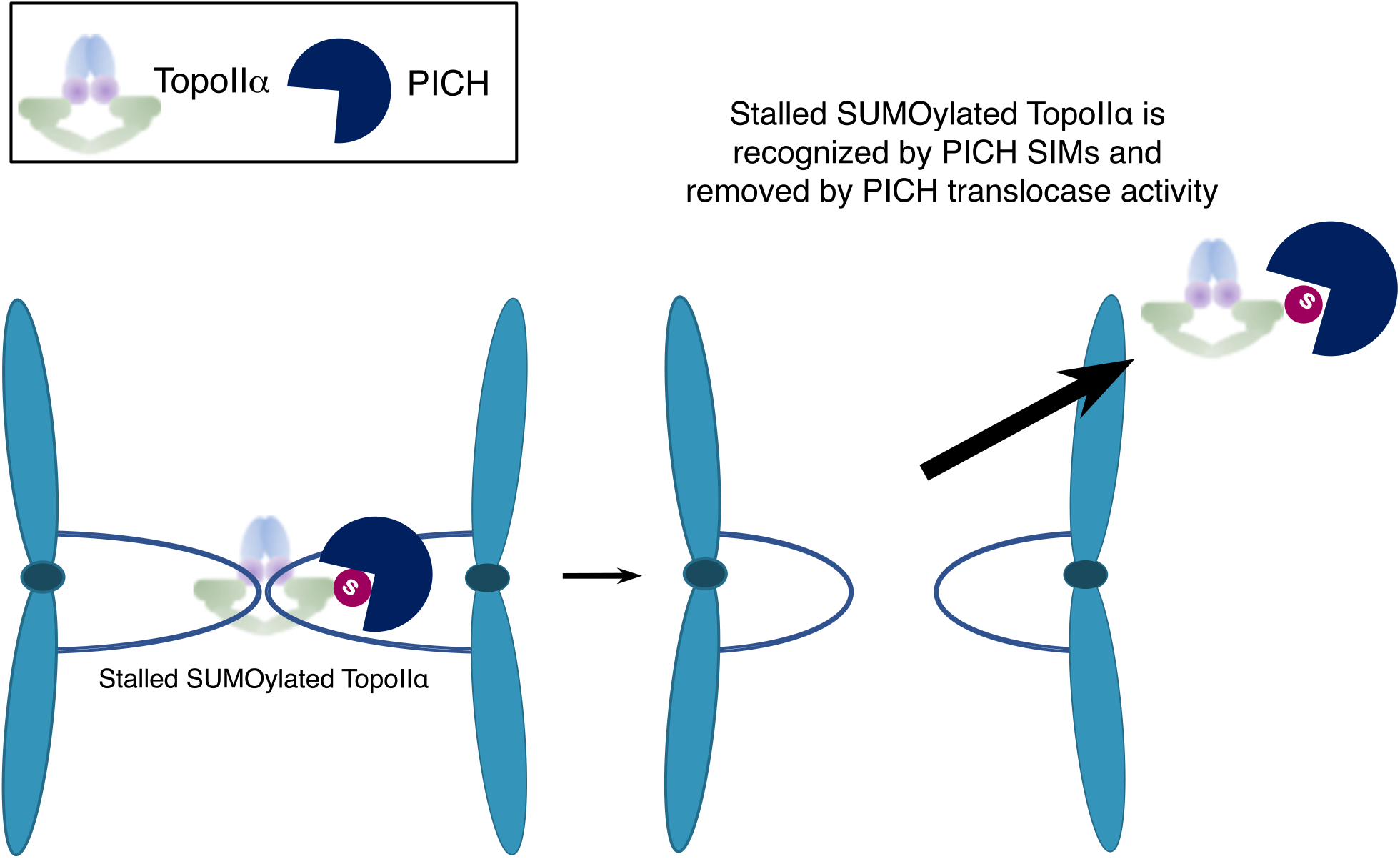
The model for exhibiting the role of PICH on stalled SUMOylated TopoIIα to promote sister chromatid disjunction. During the metaphase-to-anaphase transition, TopoIIα decatenates the last tangles of cohesed DNA between sister chromatids. The interlinked DNA molecules are released by the TopoIIα Strand Passage Reaction (SPR). If TopoIIα is stalled during the SPR, decatenated DNA molecules are bound within TopoIIα protein, and this ultimately results in the formation of chromosome bridges. This TopoIIα conformation is particularly susceptible to SUMOylation, thus becomes a critical target of PICH through its SIMs. WT PICH binds to SUMOylated TopoIIα and removes stalled SUMOylated TopoIIα using its translocase activity. Once the SUMOylated TopoIIα is removed from DNA, the sister chromatids undergo faithful segregation. In contrast, if SUMOylation is attenuated, or PICH-SIMs are mutated, chromosome bridges will form because SUMOylated TopoIIα will not be removed. If the ATPase domain of PICH is mutated, it still binds to the SUMOylated TopoIIα, however, it does not remove TopoIIα from DNA due to the loss of translocase activity, thus results in chromosome bridge formation.

## Discussion

The identification of PICH led to the discovery of UFBs which represent the existence of tangled DNA during mitosis (Biebricher et al., 2013, Wang et al., 2008). The importance of TopoIIα in resolving UFBs is highlighted by a study showing an increased incidence of PICH-positive UFBs in TopoIIα-knockdown cells (Spence et al., 2007). Likewise, knocking out PICH sensitizes cells to ICRF-193 treatment, suggesting that PICH plays a role in resolving stalled TopoIIα mediated UFB formation (Kurasawa and Yu-Lee, 2010, Nielsen et al., 2015). The current model indicates that the requirement of PICH in ICRF-193 treated cells is due to the necessity of PICH to increase TopoIIα decatenation activity. (Nielsen et al., 2015). However, ICRF-193 causes TopoII to stall at the last stage of the SPR when two DNA strands are held within TopoII. Thus, increasing the activity of TopoIIα by PICH does not entirely explain how this would lead to the resolution of stalled TopoIIα. Therefore, we propose an advanced model showing how PICH directly removes stalled SUMOylated TopoIIα from chromosomes in the presence of ICRF-193 (Figure 8). This model is supported by conditional knockdown of PICH that showed increased retention of SUMOylated TopoIIα on mitotic chromosomes (Figure 4). By treating the ΔPICH cells with ICRF-193, the retention of SUMOylated TopoIIα became more significant, supporting the specific role of PICH in removing stalled SUMOylated TopoIIα. *In vitro* assays further support that PICH utilizes its SIMs and its translocase activity to attenuate SUMOylated TopoIIα decatenation activity (Figure 7C, D). ICRF-193 stalls TopoIIα in a closed clamp conformation with two DNA strands are bound within it, and this structure is particularly susceptible to SUMOylation. PICH then binds SUMOylated TopoIIα utilizing its SIMs and removes it from its stalled position using its translocase activity, resulting in the release of two resolved DNA strands held by stalled TopoIIα (Figure 8). The process of removing stalled SUMOylated TopoIIα from decatenated, but not released DNA, resolves chromosome bridges which were originally shown to be upregulated in the PICH knockout/knockdown experiments (Kurasawa and Yu-Lee, 2010, Nielsen et al., 2015).

One remaining question is how SUMOylated TopoIIα becomes a critical target of PICH among all of the SUMOylated chromosomal proteins in the ICRF-193 treated cells. Our *in vitro* assays and previous reports showed that PICH interacts with TopoIIα and affects TopoIIα activity (Nielsen et al., 2015). This suggests that PICH has a binding affinity for TopoIIα regardless of its modification status, thus due to this intrinsic binding affinity PICH preferentially binds SUMOylated TopoIIα over other SUMOylated proteins. Another possibility is the contribution of other posttranslational modifications on TopoIIα that are influenced by ICRF-193 treatment. TopoIIα is known to be phosphorylated at its C-terminal domain with ICRF-193 treatment in mammalian cells and fission yeast (Luo et al., 2009, Nakazawa et al., 2019). The phosphorylation is suggested to play a critical role in a TopoII-dependent cell cycle checkpoint. We demonstrated that SUMOylation of TopoIIα promotes binding with Claspin (Ryu et al., 2015) which is an upstream regulator of Chk1 (Kumagai and Dunphy, 2003), and Haspin (Yoshida et al., 2016) the kinase responsible for phosphorylating H3T3 (Dai et al., 2005). In the future, it is essential to study whether kinases bound to SUMOylated TopoIIα affect the phosphorylation of TopoIIα and the activity of PICH for removing SUMOylated TopoIIα from chromosomes.

Although SUMOylated TopoIIα is a critical target of PICH in the ICRF-193 treated cells, PICH also interacts with other SUMOylated proteins and may control their binding to chromosomes. This is supported by our results which show retention of other SUMO2/3 modified proteins on mitotic chromosomes in ΔPICH cells (Supplemental Figure 6B +Auxin lane). We previously showed that PICH interacts with SUMOylated PARP1 as well as a tetrameric SUMO chain, suggesting that PICH promiscuously binds SUMOylated proteins (Sridharan et al., 2015). Both loss of translocase activity and SUMO-binding activity of PICH leads to chromosome bridge formation (Sridharan and Azuma, 2016) which could derive from the increased incidence of UFBs due to stalled SUMOylated TopoIIα. However, other SUMO2/3 modified chromosomal proteins remodeled by PICH might contribute to chromosome bridge formation in loss of PICH cells (Baumann et al., 2007, Kurasawa and Yu-Lee, 2010, Nielsen et al., 2015). Supporting this idea, it has been shown that defects in the regulation of mitotic SUMOylation causes similar chromosome bridge formation. For example, loss of a SUMO E3 ligase showed mitotic defects and chromosome bridge formation in *Drosophila* (Hari et al., 2001). Also, defects in deSUMOylation enzymes induce defective mitosis with chromosome bridge formation in cultured cells (Cubenas-Potts et al., 2013, Mukhopadhyay et al., 2010, Zhang et al., 2008). Several potential key SUMOylated chromosomal proteins were proposed to explain this SUMOylation-dependent defect (Myatt et al., 2014, Schimmel et al., 2014, Zhang et al., 2008). Once we identify which SUMOylated chromosomal proteins are controlled by PICH, and characterize their abundance on chromosomes, we will be able to elucidate the role of PICH as a “SUMOylated chromosomal protein remodeler” and its comprehensive function in chromosome segregation.

## Materials and Methods

### Plasmids, constructs, and site-directed mutagenesis

The Py-S2 fusion DNA construct of human PIASy-NTD (amino acid 1-135) and SENP2-CD (amino acid 363-589) was created by fusion PCR method using a GA linker between the two fragments. Then, the Py-S2 fusion DNA fragment was subcloned into a recombinant expression pET28a plasmid at the BamHI/XhoI sites. To generate the Py-S2 Mut fusion DNA construct, substitution of Cysteine to Alanine at 548 in Py-S2 was introduced using a site-directed mutagenesis QuikChangeII kit (Agilent) by following the manufacturer’s instructions. AAVS1 locus targeting donor plasmids for inducible expression of Py-S2 proteins were created by modifying pMK243 (Tet-OsTIR1-PURO) plasmid (Natsume et al., 2016). pMK243 (Tet-OsTIR1-PURO) was purchased from Addgene (#72835) and the OsTIR1 fragment was removed by BglII and MluI digestion, followed by an insertion of a multi-cloning site. The Py-S2 fragments were inserted at the MluI and SalI sites of the modified pMK243 plasmid. The original plasmid for OsTIR1 targeting to RCC1 locus was created by inserting the TIR1 sequence amplified from pBABE TIR1-9Myc (Addgene #47328; (Holland et al., 2012) plasmid, Blasticidin resistant gene (BSD) amplified from pQCXIB with ires-blast (Takara/Clontech), and miRFP670 amplified from pmiRFP670-N1 plasmid (Addgene #79987; (Shcherbakova et al., 2016) into the pEGFP-N1 vector (Takara/Clontech) with homology arms for RCC1 C-terminal locus. Using genomic DNA obtained from DLD-1 cell as a template DNA, the homology arms were amplified using primers listed in supplemental information (Supporting information Table 1). Further, OsTIR1 targeting plasmid was modified by eliminating the miRFP670 sequence by PCR amplification of left homology arm and TIR/BSD/right homology arm for inserting into pMK292 obtained from Addgene (#72830) (Natsume et al., 2016) using XmaI/BstBI sites. Three copies of codon optimized micro AID tag (50 amino-acid each (Morawska and Ulrich, 2013)) was synthesized by the IDT company, and hygromycin resistant gene/ P2A sequence was inserted upstream of the 3x micro AID sequence. The 3xFlag sequence from p3xFLAG-CMV-7.1 plasmid (Sigma) was inserted downstream of the AID sequence. The homology arms sequences for PICH N-terminal insertion and TopoIIα N-terminal insertion were amplified using primers listed in supplemental information (Table S1) from genomic DNA of DLD-1 cell, then inserted into the plasmid by using PciI/SalI and SpeI/NotI sites. In all of RCC1 locus, PICH locus, and TopoIIα locus genome editing cases, the guide RNA sequences listed in supplemental information (Table S1) were designed using CRISPR Design Tools from https://figshare.com/articles/CRISPR_Design_Tool/1117899 (Rafael Casellas laboratory, NIH) and http://crispr.mit.edu:8079 (Zhang laboratory, MIT) inserted into pX330 (Addgene #42230) vector using the Zhang Lab General Cloning Protocol (Cong et al., 2013). Guide plasmid for targeting AAVS1 locus (AAVS1 T2 CRIPR in pX330) was obtained from Addgene (#72833) (Natsume et al., 2016). Mutations were introduced in PAM sequences on the homology arms. The *X. laevis* TopoIIα cDNA and human PICH cDNA were subcloned into a pPIC 3.5K vector in which calmodulin-binding protein CBP-T7 tag sequences were inserted as previously described (Ryu et al., 2010b, Sridharan and Azuma, 2016). All mutations in the plasmids were generated by site-directed mutagenesis using a QuikChangeII kit (Agilent) according to manufacturer’s instructions. All constructs were verified by DNA sequencing.

### Recombinant protein expression and purification, and preparation of antibodies

Recombinant TopoIIα and PICH proteins were prepared as previously described (Ryu et al., 2010b, Sridharan and Azuma, 2016). In brief, the pPIC 3.5K plasmids carrying TopoIIα or PICH cDNA fused with Calmodulin binding protein-tag were transformed into the GS115 strain of *Pichia pastoris* yeast and expressed by following the manufacturer’s instructions (Thermo/Fisher). Yeast cells expressing recombinant proteins were frozen and ground with coffee grinder that contain dry ice, suspended with lysis buffer (50 mM Tris-HCl, pH 7.5, 150 mM NaCl, 2 mM CaCl_2_, 1 mM MgCl_2_, 0.1% Triton X-100, 5% glycerol, 1 mM DTT, complete EDTA-free Protease inhibitor tablet (Roche), and 10 mM PMSF). The lysed samples were centrifuged at 25,000 *g* for 40 min. To capture the CBP-tagged proteins, the supernatant was mixed with calmodulin-sepharose resin (GE Healthcare) for 90 min at 4°C. The resin was then washed with lysis buffer, and proteins were eluted with buffer containing 10 mM EGTA. In the case of PICH, the elution was concentrated by centrifugal concentrator (Amicon ultra with a 100kDa molecular weight cut-off). In the case of TopoIIα, the elution was further purified by Hi-trap Q anion-exchange chromatography (GE Healthcare). Recombinant Py-S2 proteins fused to hexa-histidine tag were expressed in Rossetta2 (DE3) (EMD Millipore/Novagen) and purified with hexa-histidine affinity resin (Talon beads from Takara/Clontech). Fractions by imidazole-elution were subjected to Hi-trap SP cation-exchange chromatography. The peak fractions were pooled then concentrated by centrifugal concentrator (Amicon ultra with a 30kDa molecular weight cut-off). The E1 complex (Aos1/Uba2 heterodimer), PIASy, Ubc9, dnUbc9, and SUMO paralogues were expressed in Rosetta2(DE3) and purified as described previously (Ryu et al., 2010a).

To generate the antibody for human PICH, the 3’end (coding for amino acids 947~1250) was amplified from PICH cDNA by PCR. The amplified fragment was subcloned into pET28a vector (EMD Millipore/Novagen) then the sequence was verified by DNA sequencing. The recombinant protein was expressed in Rossetta2(DE3) strain (EMD Millipore/Novagen). Expressed fragment was found in inclusion body thus the proteins were solubilized by 8M urea containing buffer (20mM Hepes pH7.8, 300mM NaCl, 1mM MgCl2, 0.5mM TCEP). The solubilized fragment was purified by Talon-resin (Clontech/Takara) using the hexa-histidine-tag fused at the N-terminus of the fragment. The purified fragment was separated by SDS-PAGE and protein was excised after Instant*Blue*TM (Sigma-Aldrich) staining. The gel slice was used as an antigen and immunization of rabbits was made by Pacific Immunology Inc., CA, USA. To generate the primary antibody for human TopoIIα, the 3’end of TopoIIα (coding for amino acids 1359~1589) was amplified from TopoIIα cDNA by PCR. The amplified fragment was subcloned into pET28a and pGEX-4T vectors (GE Healthcare) then the sequence was verified by DNA sequencing. The recombinant protein was expressed in Rossetta2(DE3). The expressed peptide was purified using hexa-histidine-tag and GST-tag by Talon-resin (Clontech/Takara) or Glutathione-sepharose (GE healthcare) following the manufacture’s protocol. The purified peptides were further separated by cation-exchange column. Purified hexa-histidine-tagged TopoIIα peptide as used as an antigen and immunization of rabbits was made by Pacific Immunology Inc., CA, USA. For both PICH and TopoIIα antigens, antigen affinity columns were prepared by conjugating purified antigens (hexa-histidine-tagged PICH C-terminus fragment or GST-tagged TopoIIα C-terminus fragment) to the NHS-Sepharose resin following manufacture’s protocol (GE healthcare). The rabbit antisera were subjected to affinity purification using antigen affinity columns. Secondary antibodies used for this study and their dilution rates were: for Western blotting; Goat anti-Rabbit (IRDye_®_680RD, 1/20000, LI-COR) and Goat anti-Mouse (IRDye_®_800CW, 1/20000, LI-COR), and for immunofluorescence staining; Goat anti-mouse IgG Alexa Fluor 568 (#A11031, 1:500, Invitrogen), goat anti-rabbit IgG Alexa Fluor 568 (#A11036, 1:500, Invitrogen), goat anti-rabbit IgG Alexa Fluor 488 (#A11034, 1:500, Invitrogen), goat anti-guinea pig IgG Alexa Fluor 647 (#A21450, 1:500, Invitrogen). Unless otherwise stated, all chemicals were obtained from Sigma-Aldrich.

### *In vitro* SUMOylation assays and decatenation assays

The SUMOylation reactions performed in the Reaction buffer (20 mM Hepes, pH 7.8, 100 mM NaCl, 5 mM MgCl_2_, 0.05% Tween 20, 5% glycerol, 2.5mM ATP, and 1 mM DTT) by adding 15 nM E1, 15 nM Ubc9, 45 nM PIASy, 500 nM T7-tagged TopoIIα, and 5 µM SUMO2-GG. For the non-SUMOylated TopoIIα control, 5 µM SUMO2-G mutant was used instead of SUMO2-GG. After the reaction with the incubation for one hour at 25°C, it was stopped with the addition of EDTA at a final concentration of 10mM. For the analysis of the SUMOylation profile of TopoIIα 1.5X SDS-PAGE sample buffer was added to reaction, and the samples were resolved on 8–16% Tris-HCl gradient gels (#XP08165BOX, Invitrogen) by SDS-PAGE, then analyzed by Western blotting with HRP-conjugated anti-T7 monoclonal antibody (#T3699, EMD Millipore/Novagen).

Decatenation assays were performed in the Decatenation buffer (50 mM Tris-HCl, pH 8.0, 120 mM NaCl, 5 mM MgCl_2_, 0.5 mM DTT, 30 µg BSA/ml, and 2 mM ATP) with SUMOylated TopoIIα and non-SUMOylated TopoIIαn and with 6.2 ng/µl of kDNA (TopoGEN, Inc.). The resction was performed at 25°C with the conditions indicated in each of the figures. The reactions were stopped by adding one third volume of 6X DNA dye (30% glycerol, 0.1% SDS, 10 mM EDTA, and 0.2 µg/µl bromophenol blue). The samples were loaded on a 1% agarose gel containing SYBR_TM_ Safe DNA Gel stain (#S33102, Invitrogen) with 1kb ladder (#N3232S, NEB), and electrophoresed at 100 V in TAE buffer (Tris-acetate-EDTA) until the marker dye reached the middle of the gel. The amount of kDNA remaining in the wells was measured using ImageStudio, and the percentage of decatenated DNA was calculated as (Intensity of initial kDNA [at 0 minutes incubation] – intensity of remaining catenated DNA)/Intensity of initial kDNA. Obtained percentages of catenated DNA was plotted and analyzed for the statistics by using GraphPad Prism 8 Software.

### Cell culture, Transfection, and Colony Isolation

Targeted insertion using the CRISPR/Cas9 system was used for all integration of exogenous sequences into the genome. Either HCT116 cells or DLD-1 cells were transfected with guide plasmids and donor plasmid using ViaFect_TM_ (#E4981, Promega) on 3.5cm dishes. The cells were split and re-plated on 10cm dishes at ~20% confluency, two days after, the cells were subjected to a selection process by maintaining in the medium in a presence of desired selection reagent (1μg/ml Blasticidin (#ant-bl, Invivogen), 1μg/ml puromycin (#ant-pr, Invivogen), 200μg/ml Hygromycin B Gold (#ant-hg, Invivogen)). The cells were cultured for 10 to 14 days with a selection medium, the colonies were isolated and grown in 24 well plates, and prepared Western blotting and genomic DNA samples to verify the insertion of the transgene. Specifically, for the Western blotting analysis, the cells were pelleted, 1X SDS PAGE sample buffer was added, and boiled/vortexed. Samples were separated on an 8-16% gel and then blocked with Casein and probed using the indicated antibody described in each figure legend. Signals were acquired using the LI-COR Odyssey Fc imager. To perform genomic PCR, the cells were pelleted, genomic DNA was extracted using lysis buffer (100mM Tris-HCl pH 8.0, 200mM NaCl, 5mM EDTA, 1% SDS, and 0.6mg/mL proteinase K (#P8107S, NEB)), and purified by ethanol precipitation followed by resuspention with TE buffer containing 50ug/mL RNase A (#EN0531,ThermoFisher). Primers used for confirming the proper integrations are listed in the supplemental information.

To establish AID cell lines, as an initial step, the *Oryza sativa* E3 ligase (OsTIR1) gene was inserted into the 3’ end of a housekeeping gene, RCC1, using CRISPR/Cas9 system in the DLD-1 cell line. The RCC1 locus was an appropriate locus to accomplish the modest but sufficient expression level of the OsTIR1 protein so that it would not induce a non-specific degradation without the addition of Auxin (Supplemental Figure S1). We then introduced DNA coding for AID-3xFlag tag into the TopoIIα or PICH locus using CRISPR/Cas9 editing into the OsTIR1 expressing parental line (Supplemental Figure S2A). The isolated candidate clones were subjected to genomic PCR and Western blotting analysis to validate integration of the transgene (Supplemental Figure S2 B, C and S4 B, C). Once clones were established and the transgene integration was validated, the depletion of the protein in the auxin-treated cells was confirmed by Western blotting and immunostaining (Supplemental Figure S2D and S4C).

### Xenopus egg extract assay for mitotic chromosomal SUMOylation analysis

Low speed cytostatic factor (CSF) arrested Xenopus egg extracts (XEEs) and demembraned sperm nuclei were prepared following standard protocols (Murray, 1991, Powers et al., 2001). To prepare the mitotic replicated chromosome, CSF extracts were driven into interphase by adding 0.6mM CaCl_2_. Demembraned sperm nuclei were added to interphase extract at 4000 sperm nuclei/μl, then incubated for ~60 min to complete DNA replication confirmed by the morphology of nuclei. Then, equal volume of CSF XEE was added to the reactions to induce mitosis. To confirm the activities of Py-S2 proteins on mitotic SUMOylation, the Py-S2 proteins or dnUbC9 were added to XEEs at a final concentration of 30nM and 5μM, respectively, at the onset of mitosis-induction. After mitotic chromosome formation was confirmed by microscopic analysis of condensed mitotic chromosomes, chromosomes were isolated by centrifugation using 40% glycerol cushion as previously described (Yoshida et al., 2016) then the isolated mitotic chromosomes were boiled in SDS-PAGE sample buffer. Samples were resolved on 8-16% gradient gels and subjected to Western blotting with indicated antibodies. Signals were acquired using LI-COR Odyssey Fc digital imager and the quantification was performed using Image Studio Lite software.

The following primary antibodies were used for Western blotting: Rabbit anti-Xenopus TopoIIα (1:10,000), Rabbit anti-Xenopus PARP1 (1:10,000), Rabbit anti-SUMO2/3 (1:1,000) (all prepared as described previously (Ryu et al., 2010a)), anti-Histone H3 (#14269, Cell Signaling).

### Preparation of mitotic cells and chromosome isolation

HCT116 or DLD-1 cells were grown in McCoy’s 5A 1x L-glutamine 10% FBS media for no more than 10 passages. To analyze mitotic chromosomes, cells were synchronized by Thymidine/Nocodazole cell cycle arrest protocol. In brief, cells were arrested with 2mM Thymidine for 17 hours, were released from the Thymidine block by performing three washes with non-FBS containing McCoy’s 5A 1x L-glutamine media and placed in fresh 10%FBS containing media. 6 hours after the Thymidine release, 0.1ug/mL Nocodazole was added to the cells for 4 additional hours, mitotic cells were isolated by performing a mitotic shake-off, and washed 3 times using McCoy’s non-FBS containing media to release from Nocodazole. The released cells were resuspended with 10% FBS containing fresh media and 7uM of ICRF-193, 40uM Merbarone, or equal volume DMSO, were plated on Fibronectin coated cover slips, and incubated for 20 minutes (NEUVITRO, #GG-12-1.5-Fibronectin). To isolate mitotic chromosomes, the cells were lysed with lysis buffer (250mM Sucrose, 20mM HEPES, 100mM NaCl, 1.5mM MgCl_2_, 1mM EDTA, 1mM EGTA, 0.2% TritonX-100, 1:2000 LPC (Leupeptin, Pepstatin, Chymostatin, 20mg each/ml in DMSO; Sigma-Aldrich), and 20mM Iodoacetamide (Sigma-Aldrich #I1149)) incubated for 5 minutes on ice. Lysed cells were then placed on a 40% glycerol containing 0.025% Triton-X-100 cushion, and spun at 10,000xg for 5 minutes, twice. Isolated chromosomes were then boiled with SDS-PAGE sample buffer, resolved on an 8-16% gradient gel and subjected to Western blotting with indicated antibodies. Signals of the blotting were acquired using the LI-COR Odyssey Fc machine.

The following primary antibodies were used for Western blotting: Rabbit anti-PICH (1:1,000), Rabbit anti-TopoIIα (1:20,000) (both are prepared as described above), Rabbit anti-SUMO2/3 (1:1,000), Rabbit anti-Histone H2A (1:2,000) (#18255, Abcam), Rabbit anti-Histone H3 (1:2,000) (#14269, Cell Signaling), Rabbit anti-PIASy (1:500) (as described in (Azuma et al., 2005)), Mouse anti-β-actin (1:2,000) (#A2228, Sigma-Aldrich), Mouse anti-myc (1:1,000) (#9E10, Santa Cruz), Mouse anti-β-tubulin (1:2,000) (#, Sigma-Aldrich), Mouse anti-Flag (1:1,000) (#F1804, Sigma-Aldrich).

### Cell fixation and staining

To fix the mitotic cells on fibronectin coated cover slips, cells were incubated with 4% paraformaldehyde for 10 minutes at room temperature, and subsequently washed three times with 1X PBS containing 10mM Tris-HCl to quench PFA. Following the fixation, the cells were permeabilized using 100% ice cold Methanol in −20°C freezer for 5 minutes. Cells were then blocked using 2.5% hydrolyzed gelatin for 30 minutes at room temperature. Following blocking the cells were stained with primary antibodies for 1 hour at room temperature, washed 3 times with 1X PBS containing 0.1% tween20, and incubated with secondary for 1 hour at room temperature. Following secondary incubation cells were washed 3 times with 1x PBS-T and mounted onto slide glass using VECTASHIELD_®_ Antifade Mounting Medium with DAPI (#H-1200, Vector laboratory) and sealed with nail polish. Images were acquired using the Plan Apo 100x/1.4 objective lens on a Nikon Ti Eclipse microscope equipped Exi Aqua CCD camera (Q imaging) or a Nikon TE2000-U equipped PRIME-BSI CMOS camera (Photometrics) with MetaMorph imaging software. Adobe Photoshop (CS6) software was used to process the images for signal intensities and size according to journal policy. The Fiji colocalization threshold software was used to measure colocalization coefficients for at least 20 cells in three independent experiments.

The following primary antibodies were used for staining: Rabbit anti-PICH 1:800, Rabbit anti-human TopoIIα 1:1000 (both are prepared as described above), Mouse anti-human TopoIIα 1:300 (#Ab 189342, Abcam), Mouse anti-SUMO2/3 (#12F3, Cytoskeleton Inc), and Guinea Pig anti-SUMO2/3 (1:300) (prepared as previously described (Ryu et al., 2010a)).

### Statistical analysis

All statistical analyses were performed with either 1- or 2-way ANOVA, followed by the appropriate post-hoc analyses for each of the analysis using GraphPad Prism 8 software.

### Animal use

For XEE assay, frog eggs were collected from mature female *Xenopus laevis*, and sperm was obtained from matured male *Xenopus laevis*. Animal protocol for the usage of *Xenopus laevis* was approved by University of Kansas IACUC.

## Abbreviations

TopoIIα: Topoisomerase IIα
PICH: Polo-like kinase interacting checkpoint helicase
SPR: Strand passage reaction
SUMO: Small ubiquitin-like modifier
XEE: Xenopus egg extract
CSF: Cytostatic factor
nUbc9: dominant negative E2 SUMO-conjugating enzyme
SENP: Sentrin-specific protease
PIAS: Protein inhibitor of activated STAT

## Acknowledgements

We thank Dr. M. Azuma at the University of Kansas for the use of her microscope and imaging software. We also thank Dr. D. Clarke at the University Minnesota and Dr. Yukiko Yamashita at the University of Michigan for the critical reading of the manuscript and comments on this project. This work was supported by NIH/NIGMS, GM112893 and, in part, by KUCC/CB pilot grant (KAN1000623). The establishment of AID-mediated knockdown system was supported V. Aksenova, A. Arnaoutov and M. Dasso whom are supported by the National Institute for Child Health and Human Development Intramural projects Z01 HD008954 and ZIA HD001902.

## Author Contributions

VH conducted most of the experiments, created AID fused PICH cell line, prepared figures, and drafted the manuscript. HP prepared DNA constructs for genome editing, created CRISPR/Cas9 genome edited HCT116 lines for inducible expression of de-SUMOylation enzyme and for Os-TIR1 expressing DLD-1 cell line, created AID fused TopoIIα cell lines, and performed XEE assay for validation of Py-S2 proteins. NP conducted experiments for initial validation of the genome edited cell lines expressing Py-S2. BL performed analysis in Figure 6. VA, AA, and MD established AID-mediated degradation system by optimizing Os-TIR1 integration locus and creating constructs for genome editing by CRISPR/Cas9 for that system. YA designed the study, supervised project, and wrote the manuscript.

---

**Supplemental Figure S1.**
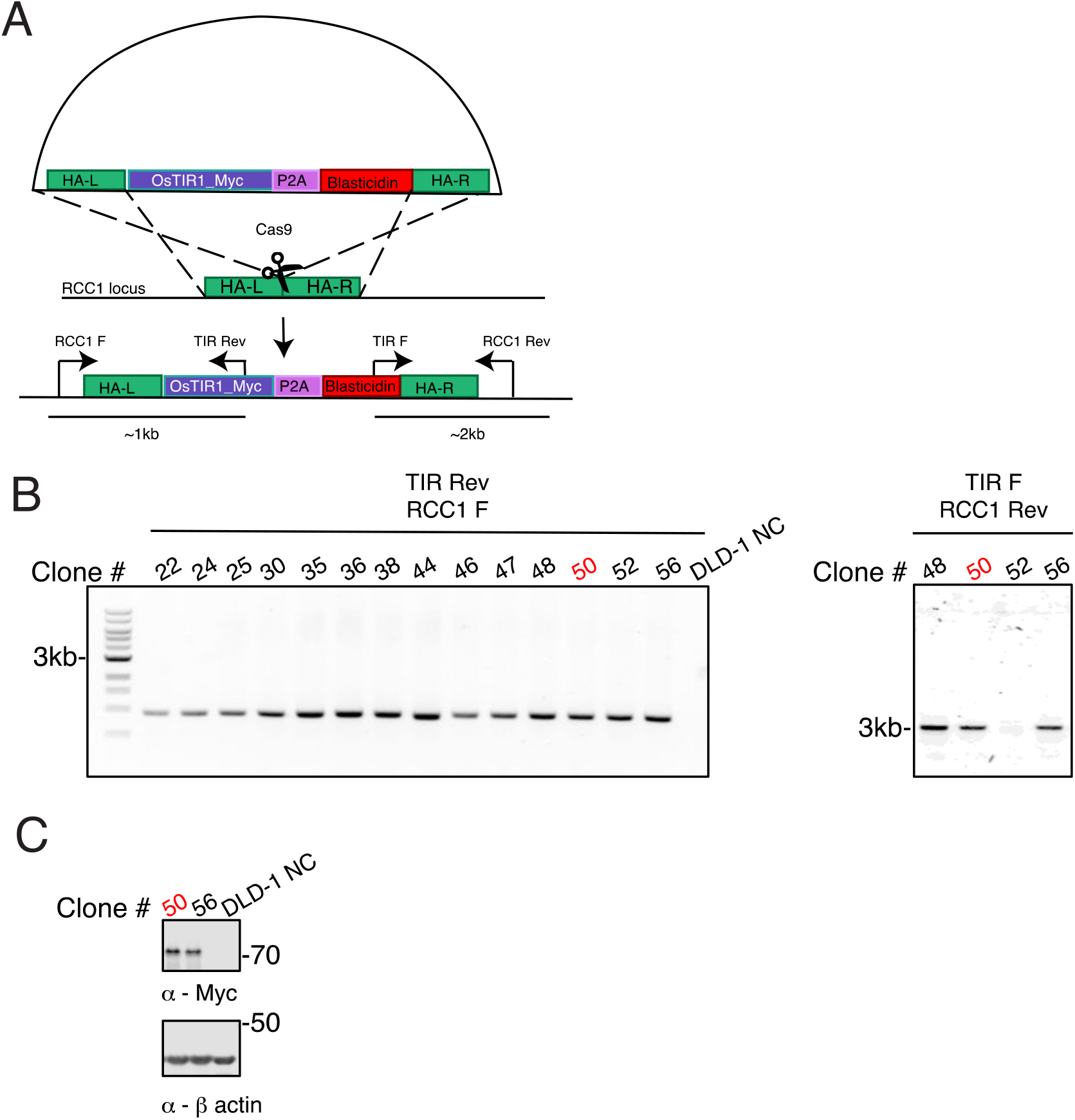
Construction of OsTIR1 expressing DLD-1 cell lines. **(A)** Experimental scheme for the establishment of OsTIR1 gene expressing DLD1 cell. RCC1-OsTIR1-Myc-P2A-Blasticidin donor plasmid, and two guide RNAs targeting the 3’ end of RCC1 were used to integrate the OsTIR1 gene into the RCC1 locus. **(B)** After the selection with 2ug/mL Blasticidin, fourteen clones were isolated and subjected to genomic PCR utilizing primers that targeted the 5’ end of the construct (upper panel). Non-transfected DLD-1 cells were used as a negative control (DLD-1 NC). Clones #48, 50, 52 and 56 were further verified by genomic PCR using primers for 3’ ends of the construct. **(C)** Among the positive clones identified in **B**, two clones were chosen to verify the protein expression by Western blotting. Whole cell lysates obtained from asynchronous cell population were subjected to Western blotting. Non-transfected DLD-1 whole cell lysate was used as a negative control (DLD-1 NC). An anti-Myc antibody was used to detect OsTIR1 protein and anti-β-actin was used as a loading control. Clone #50 (marked in red) was chosen to utilize for subsequent AID tagging for TopoIIα and PICH.

**Supplemental Figure S2.**
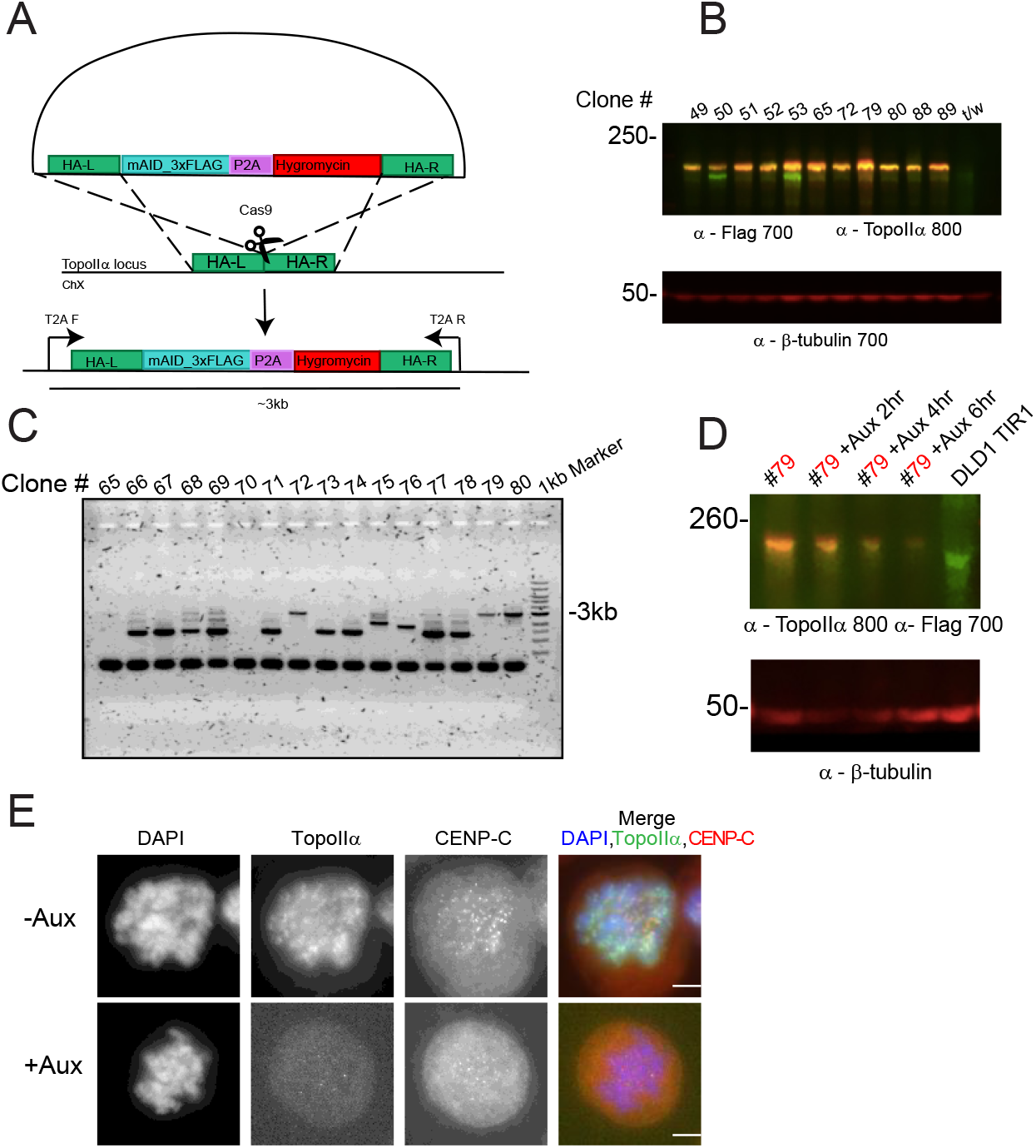
Construction of TopoIIα-AID cell line. **(A)** Experimental scheme of donor plasmid tagging the 5’ end of endogenous TopoIIα with AID. Cells were transfected with the donor plasmid together with two different guide RNAs. **(B)** After selection with 400ug/mL hygromycin, resistant clones were isolated. Whole cell lysate was obtained from cells and the expression of the transgene was screened by Western blotting analysis. Representative Western blotting of clones is shown. An anti-Flag antibody was used to detect AID-Flag tagged TopoIIα (~190kDa) in the 700 channel (red) and anti-TopoIIα antibodies were used to detect both AID-Flag tagged TopoIIα and untagged TopoIIα (~160kDa) in the 800 channel (green). Anti-β-tubulin was used as a loading control. **(C)** Genomic DNA from hygromycin resistant clones was extracted for PCR analysis using indicated primers shown in A. Representative result of PCR amplification was shown. Clones showing only 3kbp DNA fragment are homozygous AID integrated clones (#72, #79 and #80). **(D)** The clone #79 was treated with auxin for 2, 4, and 6-hours, and evaluated the TopoIIα depletion by Western blotting. As a control, DLD-1 OsTIR1#50 parental cells were treated with auxin for 6 hours (DLD1 TIR1). Whole cell lysates were subjected to Western blotting analysis using indicated antibodies. Clone #79 was chosen for further analysis in the subsequent experiments showed in Figure 2. **(E)** DLD-1 cells with endogenous TopoIIα tagged with an auxin inducible degron (AID) were synchronized in mitosis and treated with auxin 6 hours after Thymidine release. Cells were plated onto fibronectin coated coverslips and subsequently stained with anti-TopoIIα, anti-CENP-C, and DNA was labeled with DAPI. TopoIIα foci on mitotic chromosomes are completely eliminated with auxin treatment.

**Supplemental Figure S3.**
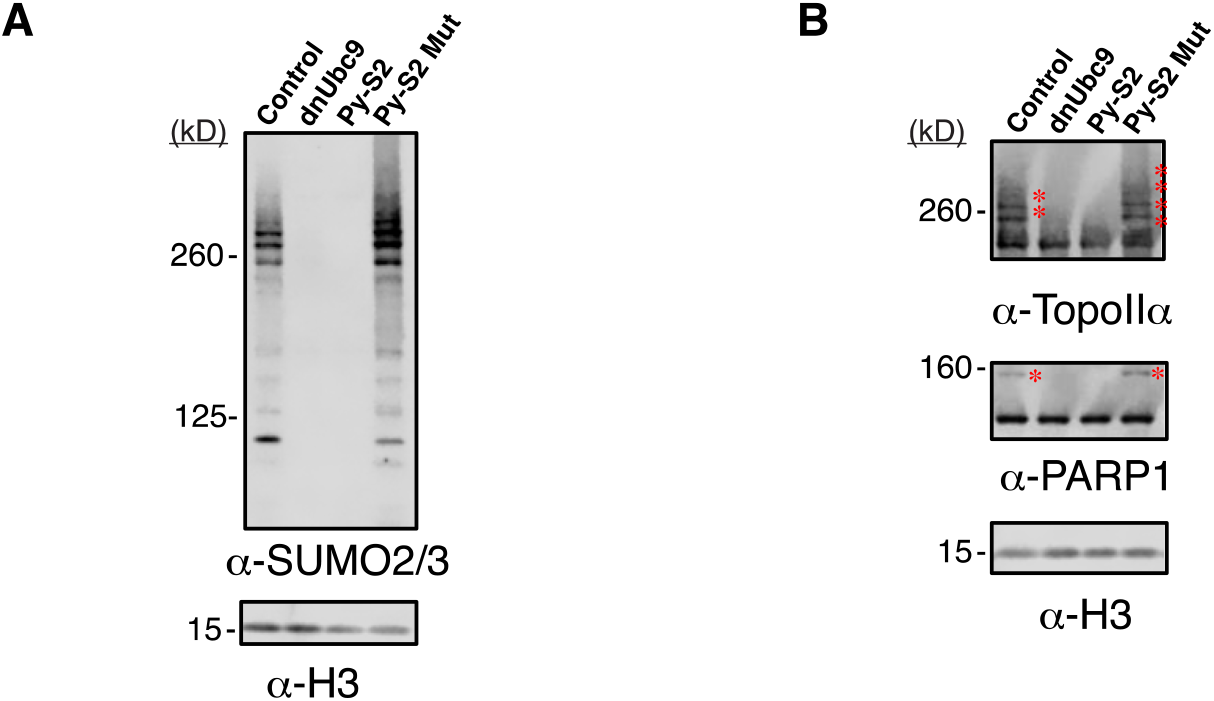
Testing SUMO modulating proteins in the *Xenopus laevis* egg extract system. **(A)** Recombinant Py-S2 or Py-S2 Mut proteins were added to *Xenopus laevis* egg extract upon induction of mitosis, and the chromosomes were isolated. Chromosome samples were subjected to Western blotting with anti-SUMO2/3 antibody. **(B)** Chromosome samples in A were subjected to Western blotting with anti-Xenopus TopoIIα antibody to detect both TopoIIα (~160kDa) and SUMOylated TopoIIα (marked with red asterisks), and anti-Xenopus PARP1 antibody to detect both PARP1 (~100kDa) and SUMOylated PARP1 (marked with red asterisks). Anti-histone H3 antibody was used as a loading control. 30nM of Py-S2 protein was sufficient to eliminate chromosomal SUMOylation, which is the equivalent concentration of endogenous PIASy protein in XEE, suggesting that the Py-S2 effectively deSUMOylates chromosomal SUMOylated proteins at a physiologically relevant concentration. Note that the concentration of dnUbc9 required for complete inhibition of chromosomal SUMOylation is 5μM in XEE, which is not within the physiological range and is difficult to induce a high expression level of dnUbc9 in cells. Addition of the Py-S2 C548A mutant (Py-S2 Mut) increased SUMO2/3 modification in chromosomal samples, including both TopoIIα SUMOylation and PARP1 SUMOylation. This suggests that the Py-S2 Mut acts as a dominant mutant for stabilizing SUMOylation.

**Supplemental Figure S4.**
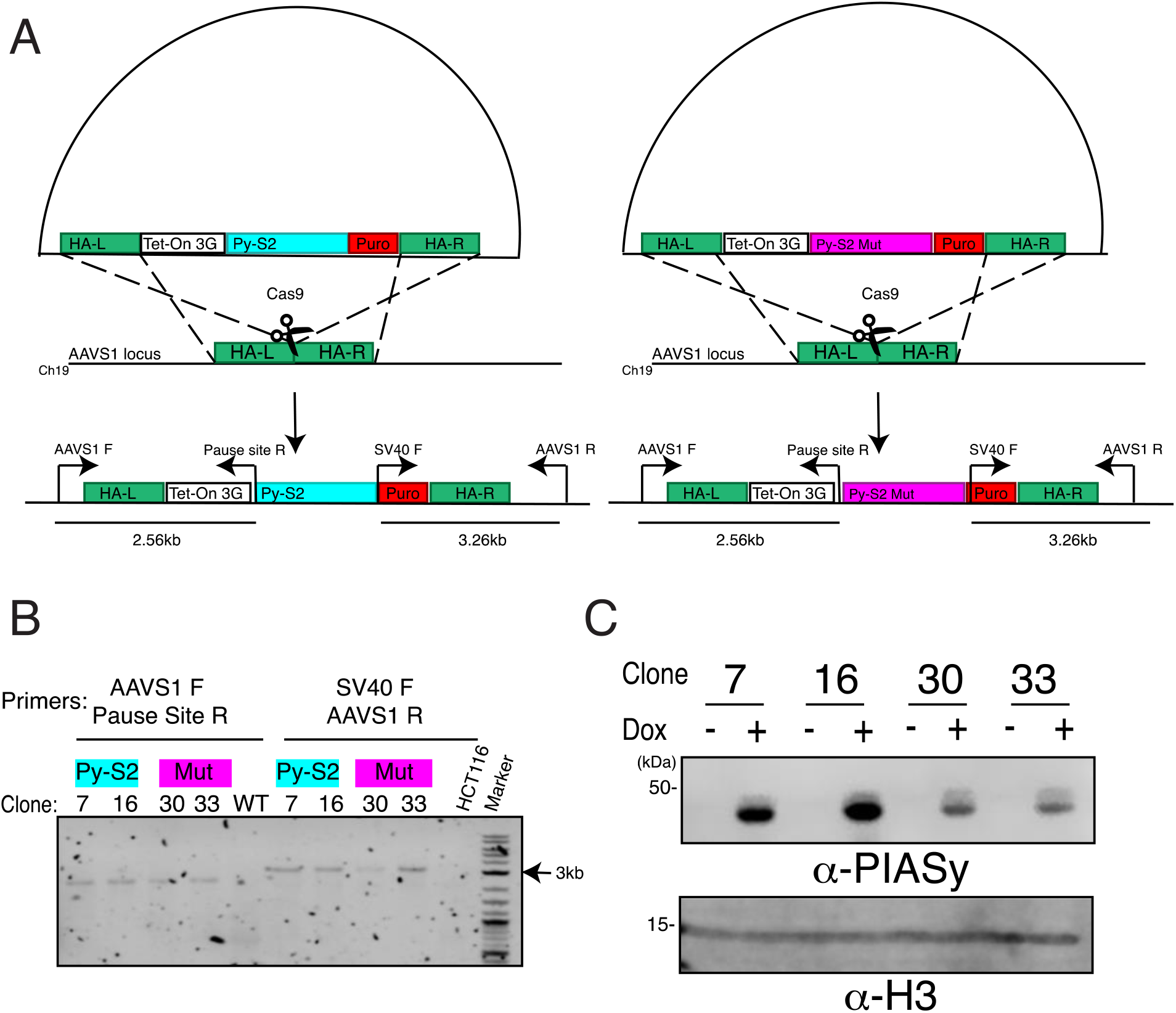
Construction of Py-S2 and Py-S2 Mut HCT116 cell lines. **(A)** Experimental scheme to introduce inducible Py-S2 and Py-S2 Mut into AAVS1 locus of HCT116 cells. Cells were transfected with a modified form of pMK243 (obtained from Addgene) AAVS1-TetON3G-Py-S2 (or Py-S2 Mut)-Puro-AAVS1 and AAVS1 T2 CRISPR/Cas9 to target AAVS1 locus. For the screening of the transgene integrated clones, primers were designed to amplify the 5’ region (2.56kb) and 3’ region (3.26kb) of the integration site respectively. **(B)** After the selection using 1ug/mL Puromycin, 2 clones each per construct were further subjected to genomic PCR to confirm the integration of the transgene. **(C)** The whole cell lysates obtained from the candidate clones were subjected to Western Blotting to confirm the inducible expression of Py-S2 and Py-S2 Mut proteins. Anti-PIASy antibodies were used to detect expression of fusion proteins (+Dox) or not (-Dox), anti-H3 antibodies were used as a loading control.

**Supplemental Figure S5.**
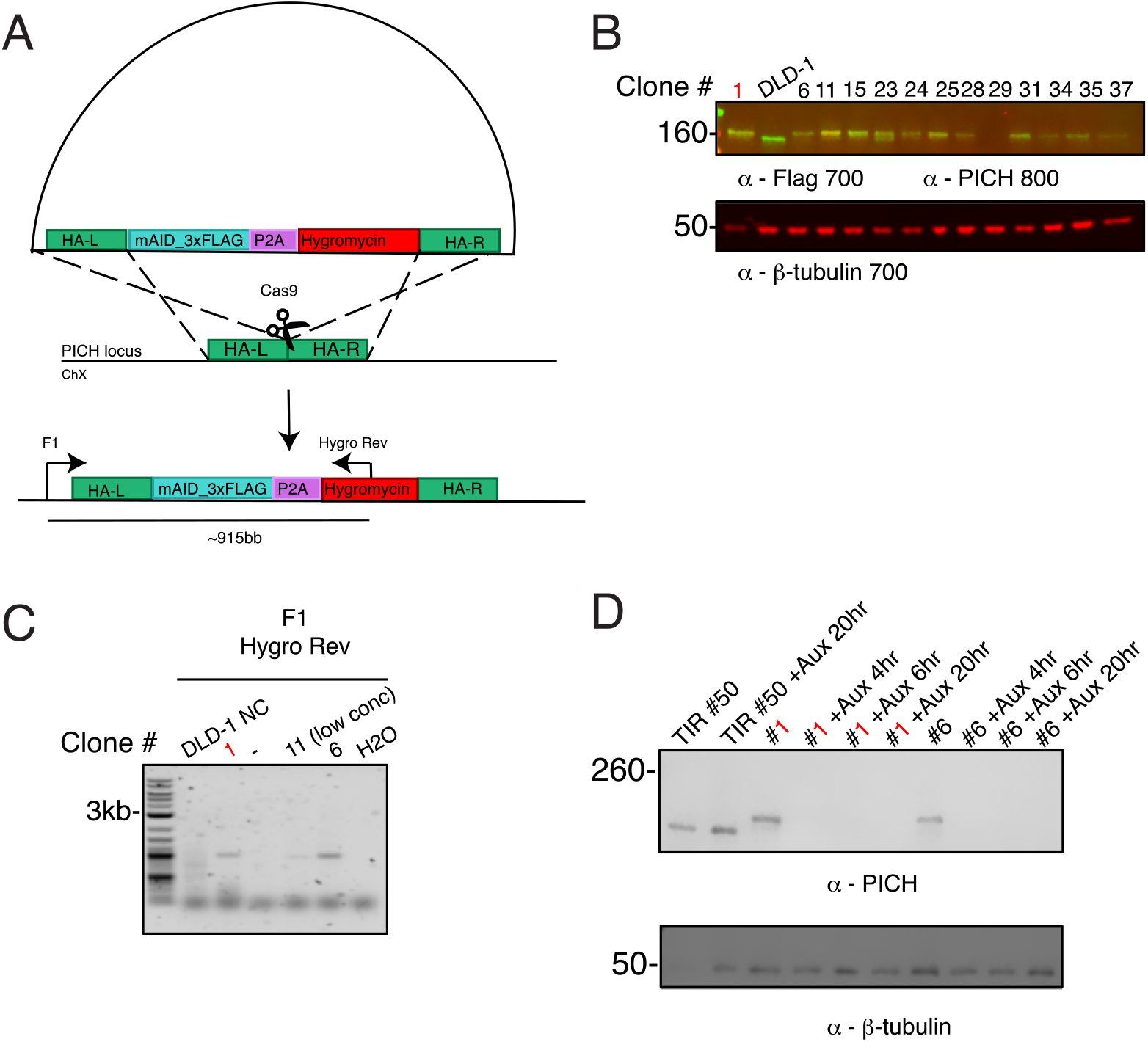
Construction of PICH-AID cell line. **(A)** Experimental scheme of donor plasmid used to tag the 5’ end of endogenous PICH locus with AID tag. Cells were transfected with PICH-mAID-3xFlag-P2A-Hygromycin donor and two different guide RNAs. After selection with 400ug/mL hygromycin clones were isolated, whole cell lysates were collected from asynchronous populations, and Western blotting was performed. **(B)** Representative Western blot for hygromycin-resistant clone screening is shown. An anti-Flag antibody was used to detect AID-Flag tagged PICH (~180kDa) in the 700 channel (colored red) and anti-PICH antibodies were used to detect both AID-Flag tagged PICH (~180kDa) and untagged PICH (~150kDa) in the 800 channel (colored green). Non-transfected DLD-1 TIR1#50 parental cell line (labeled DLD-1) was used as a negative control. Anti-β-tubulin was used as a loading control. Among thirteen samples analyzed, the clones which showed a single yellow PICH band were chosen for genomic PCR analysis (clones #1, 6 and 11). **(C)** Genomic DNA was isolated and subjected to PCR using an F1 primer located upstream of the left homology arm and Hygro Rev PCR primer located within the insert. Non-transfected DLD-1 TIR#50 parental cell DNA was used as a control (DLD-1 NC). **(D)** The clones 1 and 6 were tested for further depletion of PICH protein by auxin addition at 4, 6, and 20-hour time points. The non-transfected DLD-1 TIR1#50 parental cells were used as a control with either non-treated (TIR#50) or treated with auxin for 20 hours (TIR#50 +Aux 20 hours). The whole cell lysates were subjected to Western blotting analysis. Anti-PICH antibodies were used to detect PICH (~150kDa) or PICH-AID (~180kDa), anti-β-tubulin antibodies were used as a loading control. Clone #1 (marked in red) was chosen to utilize for subsequent experiments showed in Figure 5 and Figure S6.

**Supplemental Figure S6.**
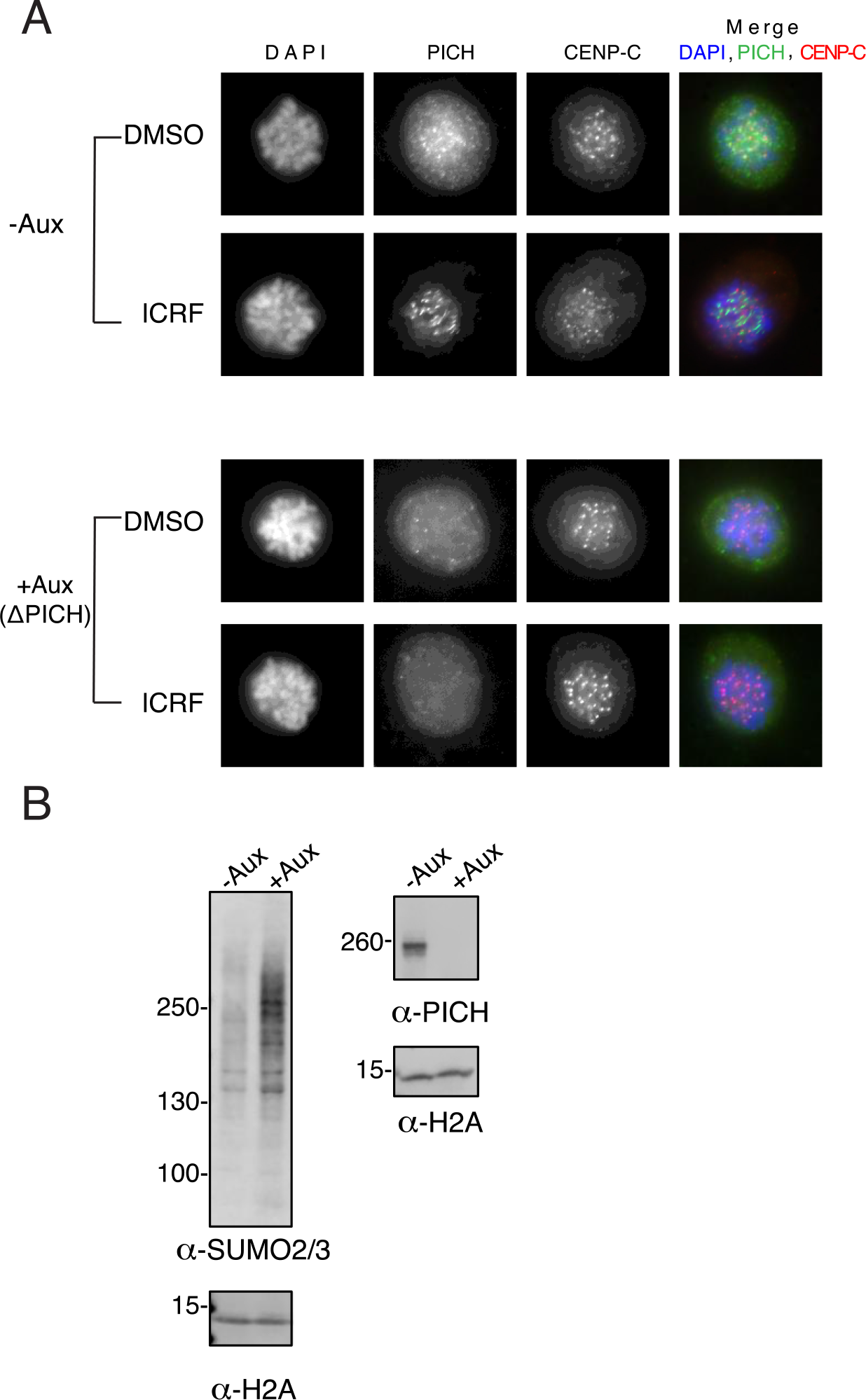
Elimination of AID-tagged PICH foci on mitotic chromosomes by addition of auxin, and effect of PICH depletion on chromosomal SUMO2/3 modified proteins. **(A)** DLD-1 cells with endogenous PICH tagged with an auxin inducible degron (AID) were synchronized in mitosis and treated with DMSO or ICRF-193. Auxin was added 6 hours after Thymidine release. Mitotic cells obtained by shake-off were plated onto fibronectin coated coverslips and subsequently stained with indicated antibodies. DNA was labeled with DAPI. PICH foci on mitotic chromosomes were completely eliminated with auxin in both DMSO and ICRF-193 treated cells. **(B)** Isolated mitotic chromosomes were subjected to Western blotting with indicated antibodies. Signals of SUMO2/3 modified chromosomal proteins are increased in +Auxin (ΔPICH) sample.

**Supporting Information Table.**
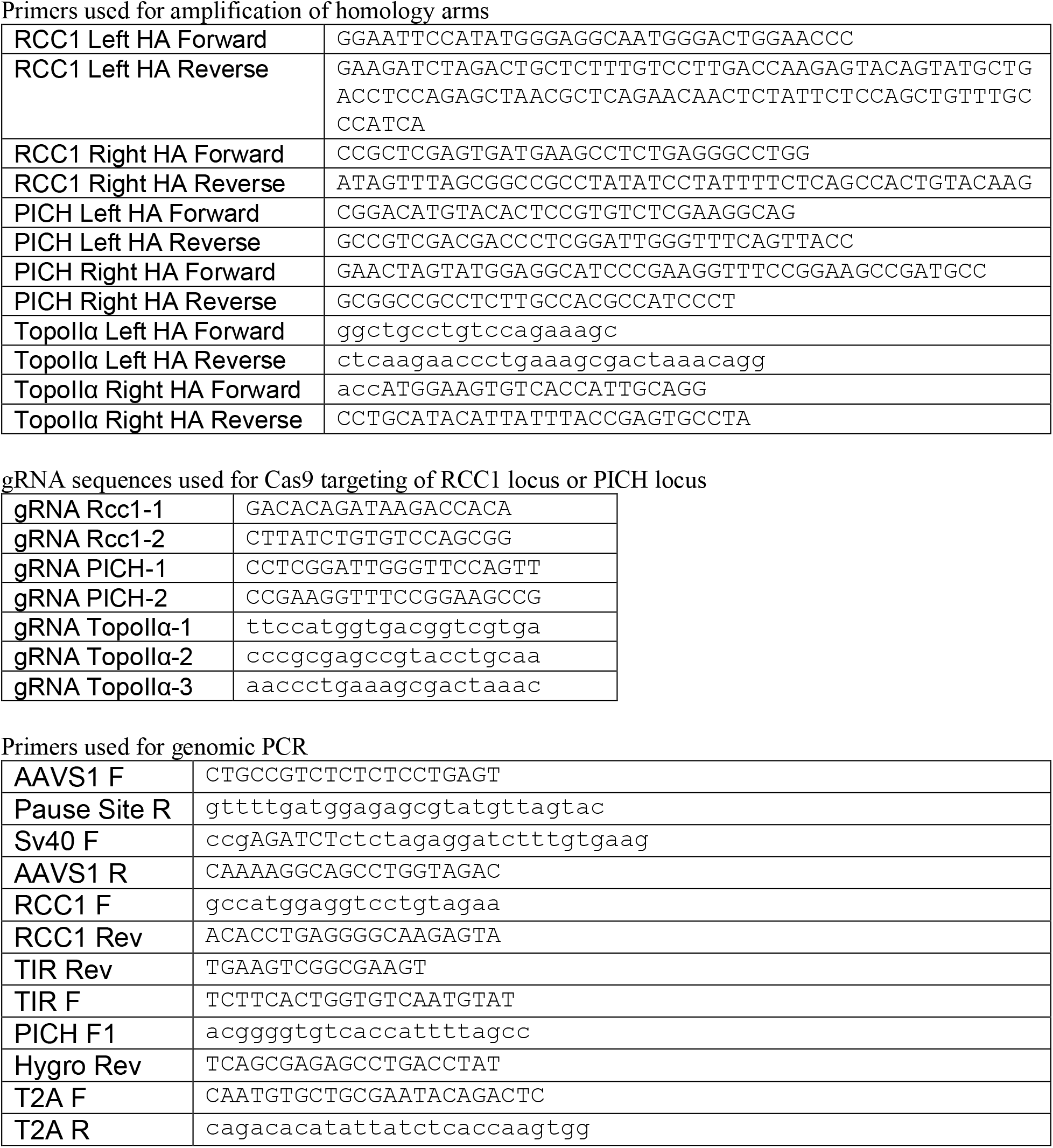

