## Supplemental figures for "PICH promotes SUMOylated TopoisomeraseIIα dissociation from mitotic centromeres for proper chromosome segregation"

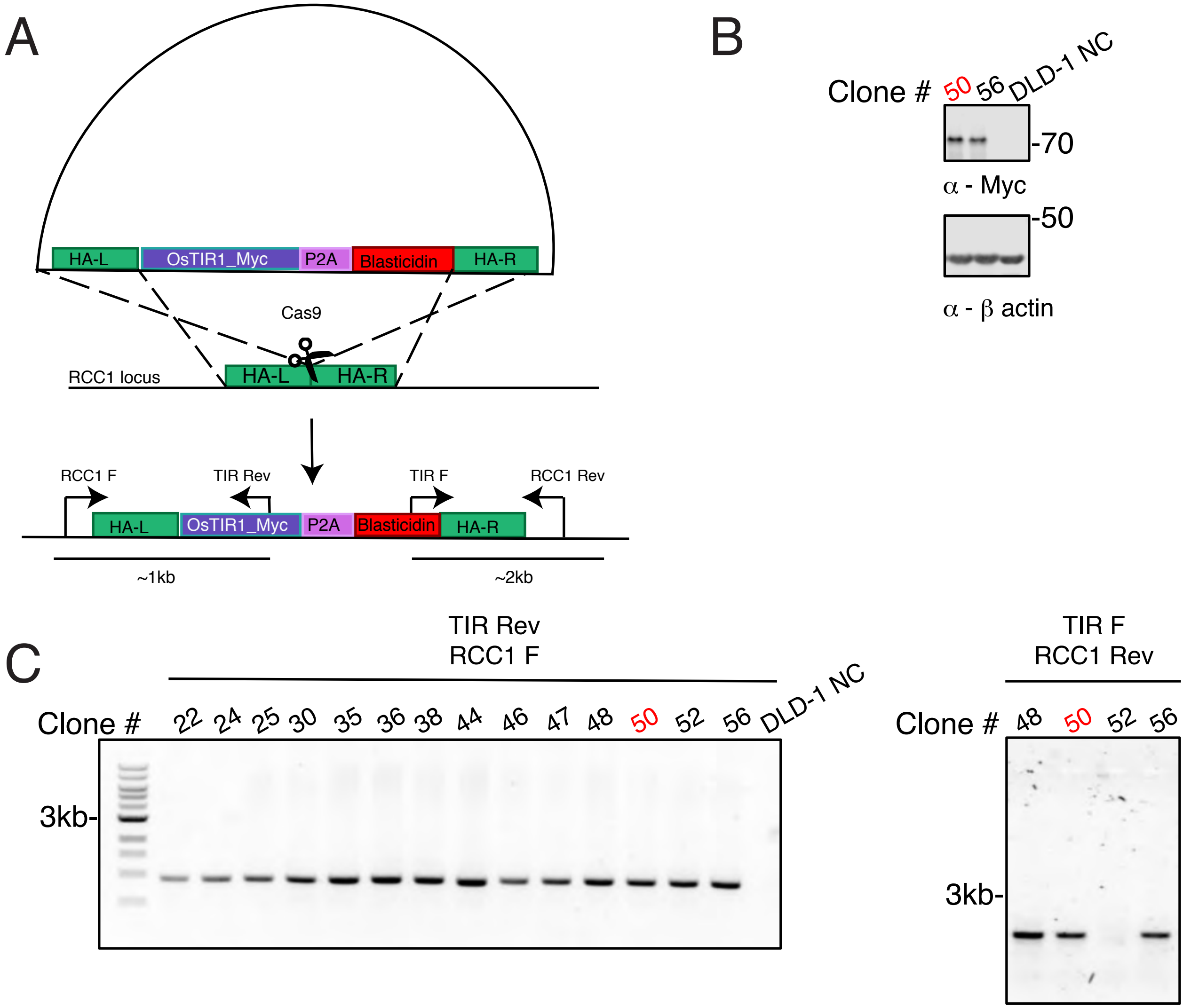


Supplemental Figure 1

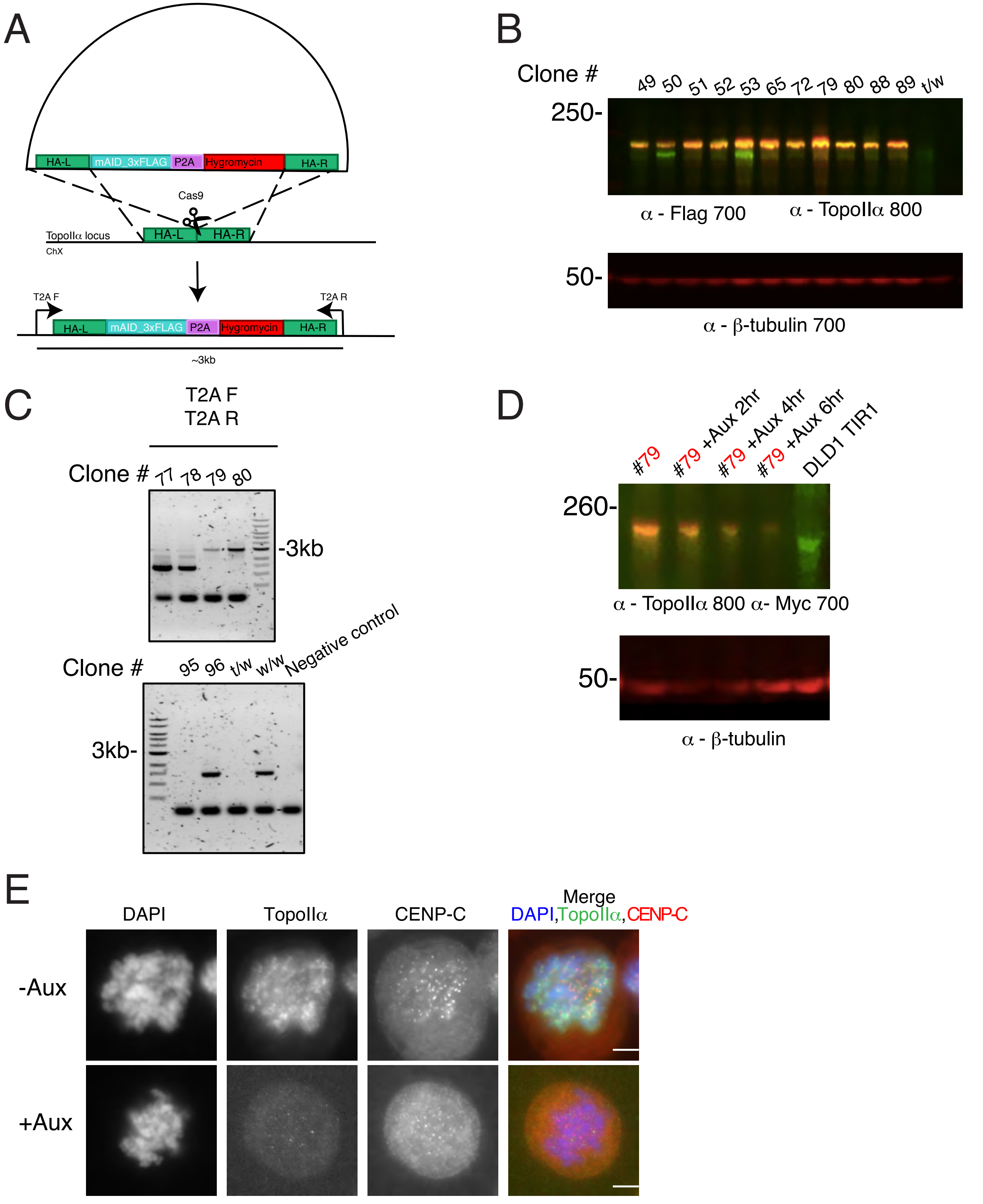


Supplemental Figure 2

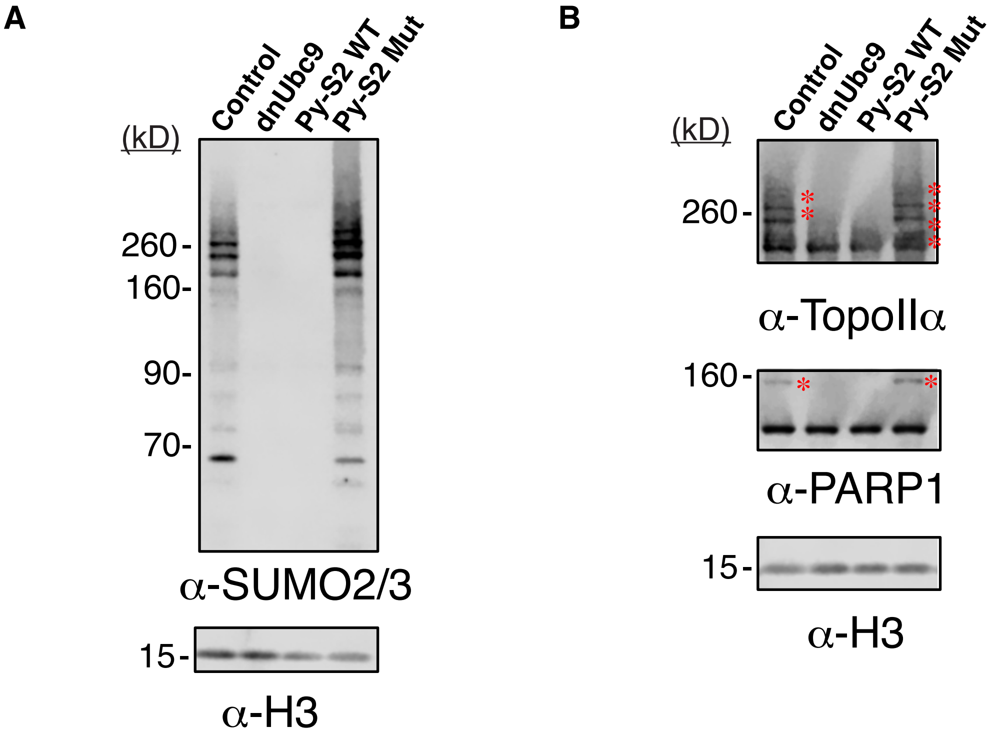


Supplemental Figure 3

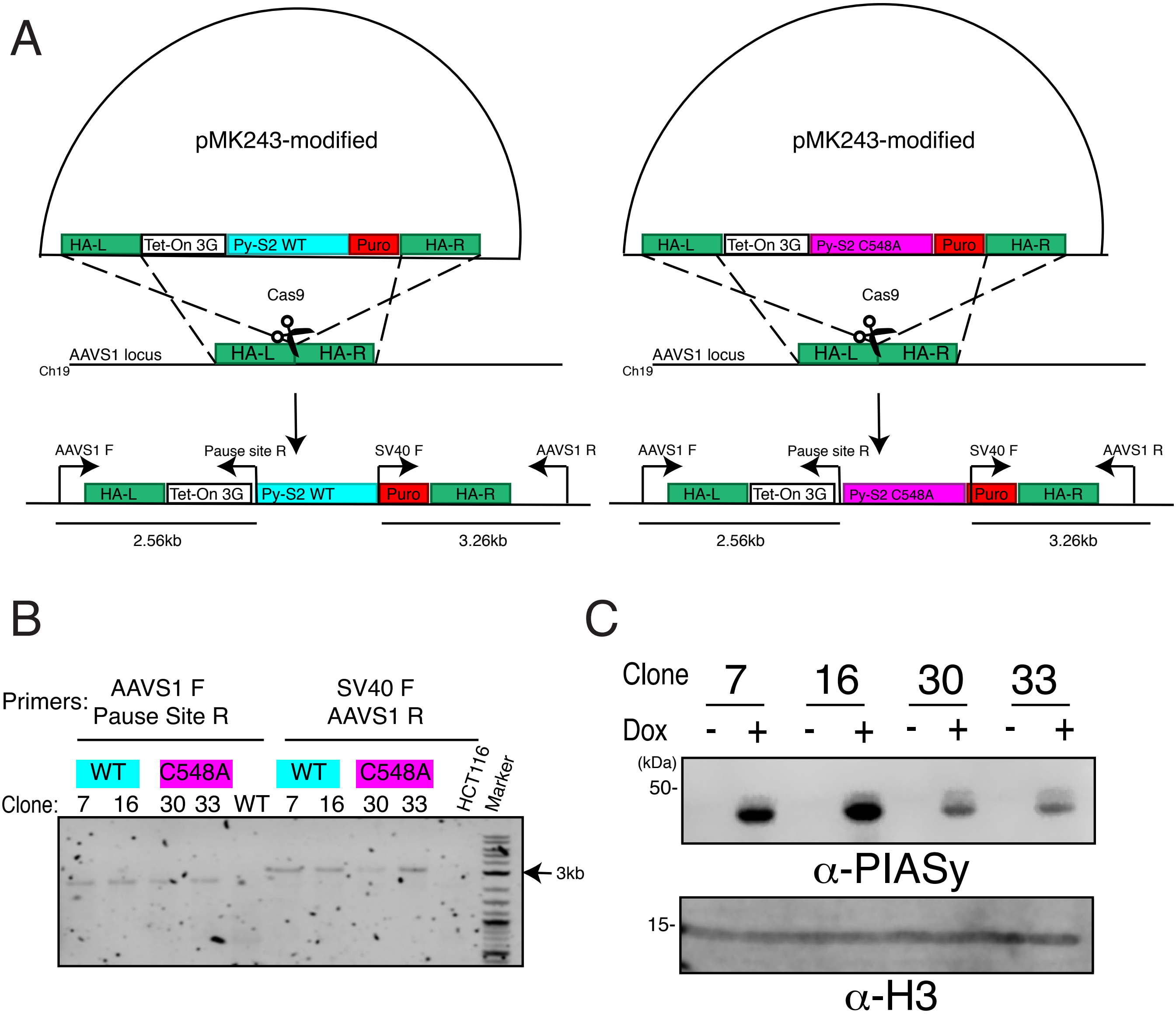


Supplemental Figure 4

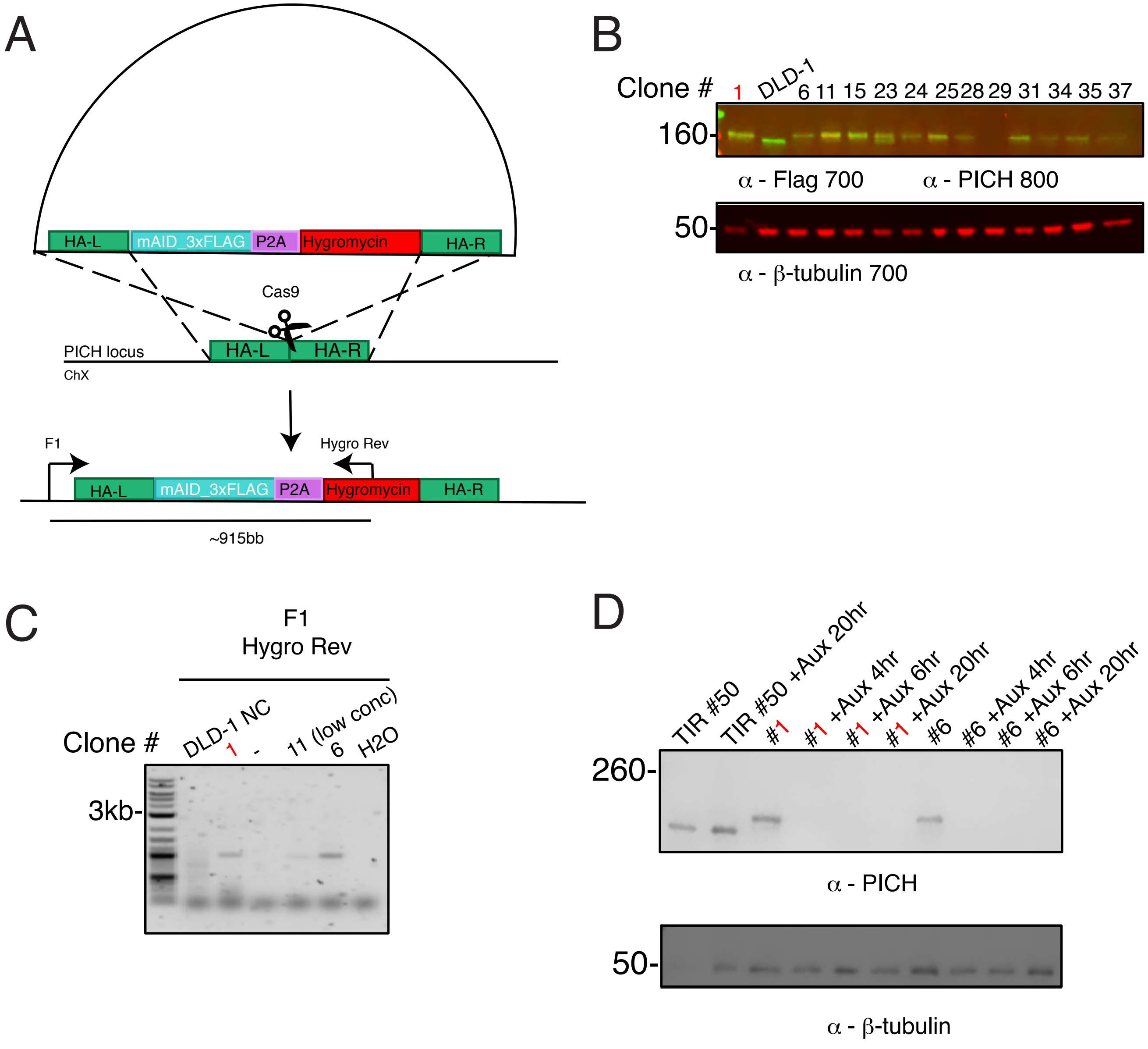


Supplemental Figure 5

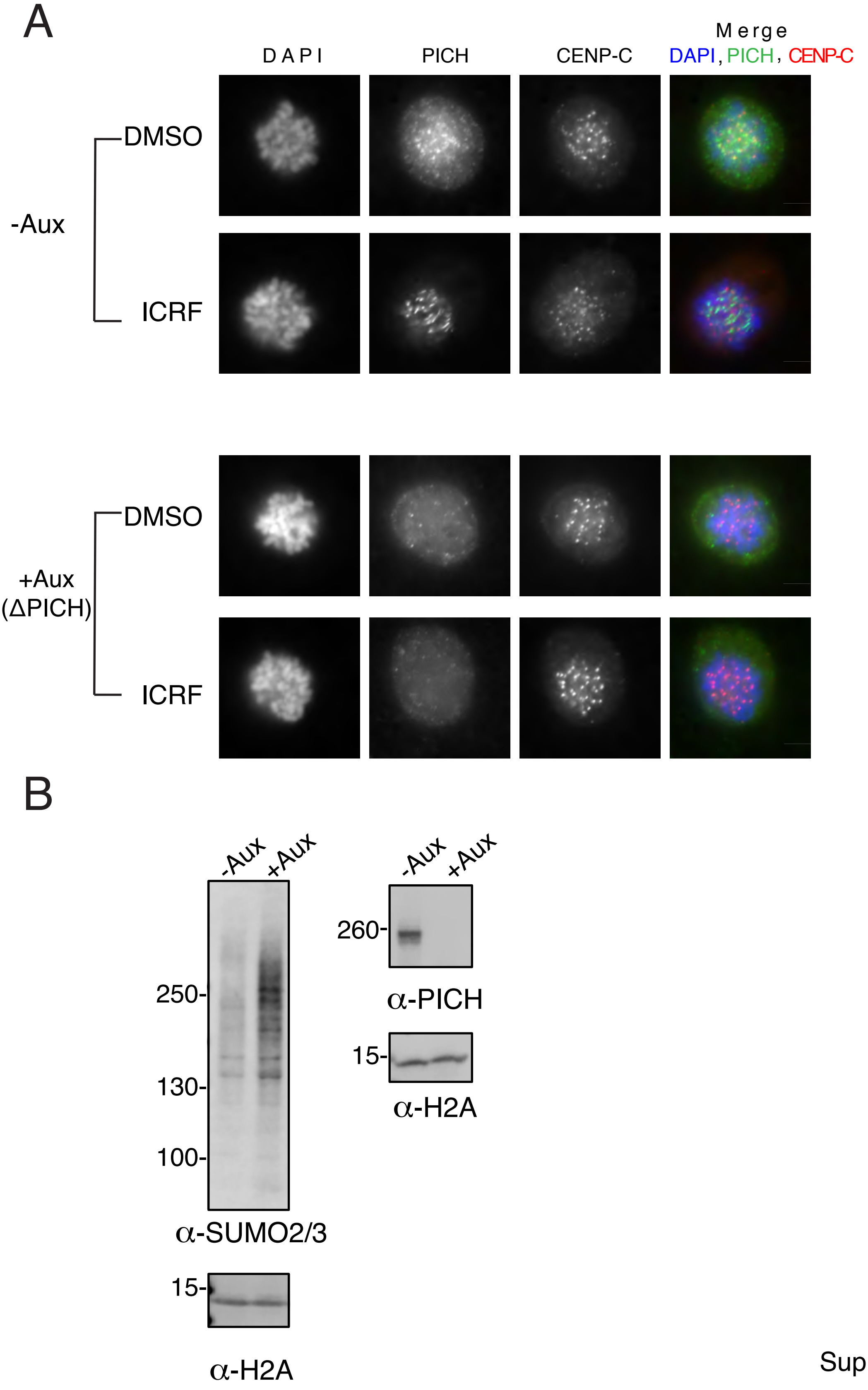


Supplemental Figure 6
